# Genomics and reproductive biology of *Leptopilina n. sp.* Buffington, Lue, Davis & Tracey sp. nov. (Hymenoptera: Figitidae): An asexual parasitoid of Caribbean *Drosophila*

**DOI:** 10.1101/2025.03.28.645512

**Authors:** Amelia RI Lindsey, Chia-Hua Lue, Jeremy S. Davis, Lydia J. Borjon, Stephanie E. Mauthner, Laura C Fricke, Lauren Eads, Molly Murphy, Melissa K Drown, Christopher Faulk, Matthew L Buffington, W Daniel Tracey

## Abstract

*Drosophila* and parasitic wasps in the genus *Leptopilina* have long been a model for understanding host-parasite interactions. Indeed, parasitic wasps are important drivers of ecological and evolutionary processes broadly, but we are generally lacking information about the diversity, natural history, and evolution of these relationships. We collected insects from the Caribbean Island of Saint Lucia, home to the eastern Caribbean ‘*dunni*’ subgroup of *Drosophila*: a clade long appreciated for its recent patterns of speciation and adaptation. Here we present an integrative approach that incorporates natural history, taxonomy, physiology, and genomics to describe *Leptopilina n. sp.* Buffington, Lue, Davis & Tracey sp. nov. (Hymenoptera: Figitidae), a virulent parasitoid of *dunni* group flies, especially *Drosophila antillea. Leptopilina n. sp.* is nested within an early-branching clade of *Leptopilina*, offering insights into the evolution of this important genus of *Drosophila* parasitoids. We present a high-quality assembly for this wasp’s 1Gbp genome, and for its bacterial endosymbiont: *Wolbachia* strain “*w*Lmal”. Furthermore, we show that *w*Lmal induces parthenogenesis in the wasp, and that these wasps are reliant upon their *Wolbachia* infections to produce female offspring. Finally, comparison to historical museum specimens indicate that *Leptopilina n. sp.* had been collected approximately 40 years prior from the nearby island of Guadeloupe and were also asexually reproducing. This work represents one of only a handful of studies in which field biology, taxonomy, systematics, genomics, and experimental biology are integrated into a species description: showcasing the possibilities for biodiversity research in the genomic era.

## INTRODUCTION

Interactions between insects and their parasitoids are ubiquitous across terrestrial ecosystems, and have been postulated as critical, but underappreciated, drivers of ecological speciation in insect groups [1, 2]. Yet compared to other ecological interactions, considerably less attention is paid to parasitism as a driver of insect speciation, in part because of the challenges of identifying often cryptic parasitoid species.

Molecular-based identification methods have rapidly revolutionized species identification, but issues persist for many parasitoid lineages [3]. Most critically, while parasitoids are commonly acknowledged as one of the most speciose groups of organisms, they are persistently under-described and under-represented in insect conservation efforts [4–6]. As such, more effort to find and describe the ecology and unique biology of new parasitoid species is needed to better understand the pattern and scope of biodiversity on earth.

Island species groups have been attractive systems for the study of speciation events and adaptive radiations since as far back as Darwin (1859), who recognized the immense number of endemic species on archipelago chains. Island archipelagos offer unique opportunities to study peripatric speciation following discrete patterns of colonization between single islands in an archipelago chain [7]. *Drosophila* species groups have been of particular interest for studying patterns of island colonization and speciation. Indeed, the Hawaiian *Drosophila* group represents one of the most well studied examples of island radiations – home to at least 500 species and research spanning decades [8–10]. While Hawaiian *Drosophila* are the most intensely studied group, several other island clades of *Drosophila* have received research attention (e.g., *sechellia-simulans* [11, 12]). Another such clade is the *D. dunni* group – a subgroup of the larger *D. cardini* group that is endemic to the Caribbean islands, particularly the Lesser Antilles [13, 14]. This subgroup is characterized by species or subspecies with distinct abdominal pigmentation endemic to just one or two major islands in this archipelago – frequently just 25-50 km apart. As such, this group represents an excellent model for the study of peripatric speciation and morphological evolution across archipelagos [12, 14]. Despite years of appreciation for the *dunni* group as a model, little is known about the parasitoid complex of these Caribbean flies.

To better understand the ecology of *dunni* group *Drosophila,* and to investigate their potential parasitoids, we (JSD, SEM, WDT) conducted an expedition to the islands of the Lesser Antilles February through May of 2019. We traveled among the islands aboard the sailing vessel Sea Salt (a 42-foot Fountaine Pajot Catamaran) which also served as a mobile laboratory. We describe here results with collected flies and associated parasitoids from the island of Saint Lucia. Other results of our expedition will be described elsewhere. We describe a new species of *Drosophila* parasitoid: *Leptopilina n. sp.* Buffington, Lue, Davis & Tracey sp. nov. (Hymenoptera: Figitidae). *Leptopilina n. sp.* appears to be endemic to the Lesser Antilles and we find that it has high virulence against *dunni* group flies, particularly *Drosophila antillea* also collected on Saint Lucia. To our knowledge, this is the first host-parasitoid association recorded for *cardini* and *dunni* group *Drosophila.* We find that the wasps reproduce via thelytokous parthenogenesis (the asexual reproduction of females) and that this is caused by infection with a bacterium: a parthenogenesis-inducing *Wolbachia* symbiont. We report *de novo* genome assemblies for *Leptopilina n. sp.* and its *Wolbachia* strain, hereinafter “*w*Lmal”. *Leptopilina n. sp.* has the largest *Leptopilina* genome reported to-date (∼1Gbp) and will serve as a useful resource for understanding the evolution of *Drosophila*-*Leptopilina* relationships in the Caribbean islands, island speciation events, and parasitoid genome evolution more broadly.

## METHODS

### Collection: Field sites and field rearing

After sailing from Wallliabou Bay of Saint Vincent and the Grenadines, we moored the research vessel and operated out of Soufrière and Marigot Bays of Saint Lucia. Verbal permission for collection of *Drosophila* and their parasitoids was obtained in a February 2019 meeting with Dr. Hannah Romain of the Saint Lucia Ministry of Agriculture. Sampling was conducted in 2019 on Saint Lucia and leveraged baited trapping with either banana or calabasa, a locally available cucurbit fruit (likely *Cucurbita moschata*).

Calabasa and banana traps contained chopped fruit mixed with water and baker’s yeast that had been kept in a closed bucket for one or more days to ferment. The Calabasa bait was chosen based on communication with Hope Hollecher who found it to be particularly attractive to *dunni* group *Drosophila.* Traps were set at two separate sites from which flies and wasps in this study derive. The first site, known as STL3 was placed on 24 February 2019 with a total of 2 calabasa traps on a short trail leading off the main road near Ladera resort and Hotel Chocolat (GPS: lat 13.83, lon –61.05, elevation 289.6m, Figure 1). The second site, known as STL6 was placed on 26 February 2019 with a total of eight traps (5 calabasa, 3 banana) along Barre d’Isle trail (GPS: lat 13.924, lon –60.959, elevation 308.7m, Figure 1). Each trapping site was checked daily for three days and insects were collected off the fruit each time. Flies were collected using either cardboard mailing tubes for vertical trapping or mouth aspiration, and wasps were collected by mouth aspiration while they searched for larvae in the bait.

**Figure 1.**
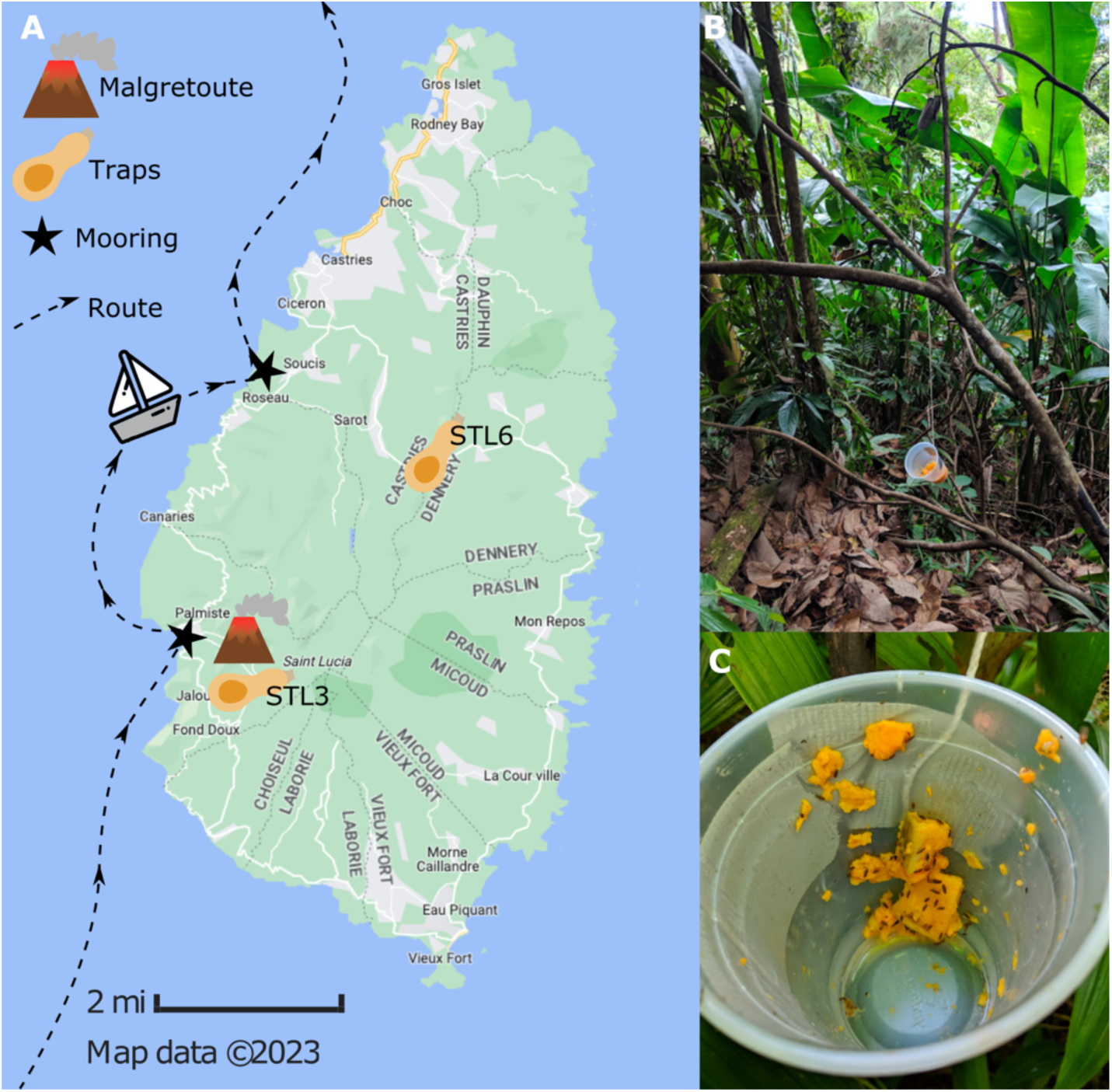
Collections on Saint Lucia: field notebook entry from JSD on 1 March 2019. “*…a 40-minute brisk hike, in which I found 0 wasps in the traps – [bummer]. Until I remembered I had set out one more trap off the path a little that I had not checked yet. And [nevertheless], my lucky [telson] finds the wasp we needed sitting in this cup. I managed to nab it and was pretty excited leaving here.*” **(A)** Map of Saint Lucia (GoogleMaps 2023), annotated with (1) locations of the STL3 and STL6 trap placements, (2) the route and mooring locations (Soufrière in the south, and Marigot in the north) of the Sea Salt, and (3) Malgretoute: the location of the La Soufrière Volcano. **(B)** The STL6 calabasa trap. **(C)** The inside of a calabasa trap.

### Collection: Colony establishment and maintenance

After collection, flies and wasps were transferred to individual rearing vials with instant fly media (Formula 4-24® Instant Drosophila Medium, Blue; Carolina Biological Supply Company). Isofemale lines of *Drosophila antillea* and *Drosophila willistoni*, were initiated using flies collected at STL3 (see results). A single wasp was collected from STL6 and was left in a vial with developing STL3 *Drosophila willistoni* larvae. This vial was then shipped back to Indiana University Bloomington under USDA APHIS Importation Permit Number P526P-19-01355. Since 2019, wasps have been maintained continuously on host cultures of *D. willistoni* on Bloomington fly food medium and further reared in IU USDA regulated containment facility number 5035.

### Taxonomy: Imaging

Voucher specimens were sent to two of us (CHL and MB) for identification. These specimens were compared with specimens deposited in the USNM as well as those on loan from Goran Nordlander (Uppsala Agricultural University, retired). Additionally, notes from Yves Carton were shared with MB, and his work with G. Nordlander in the mid 1980’s. These notes revealed an undescribed *Leptopilina* from Guadeloupe that matched the species described herein. We are therefore including these older specimens in the description. Images were acquired using a Hitachi 3000 desktop Scanning Electron Microscope (SEM), after a 120nm coating of Au-Pd from a Cressington sputter-coater. Light micrographs were shot using a Macropod 3D Pro camera setup. All type specimens collected in this study are housed in the USNM (registration numbers below); additional types, examined by G. Nordlander, reside in the Uppsala Agricultural University collection overseen by G. Nordander. Terminology for these descriptions follows Lue et al. (2016) [15].

### Taxonomy: Systematics of *Leptopilina*

Lue et al. (2016) [15] provided a detailed description on the species limits of *Leptopilina*. Species in this genus are common throughout the Holarctic Region and have recently been investigated for their biological control potential of pestiferous Drosophilidae [16, 17]. Some 29 species are currently recognized worldwide [15]. While most described species are endemic to the Holarctic Region, many undescribed species remain in collections, awaiting description. A relatively common trait among Neotropical *Leptopilina* are polychromatic antennal segments, specifically black/brown, cream, and orange-colored segments. These traits are either rare, or wholly missing, from the Paleotropics and Australia (Buffington, Lue, pers. obsv.). This is mentioned here as the new species, herein described, has rather unique color patterns among the antennal segments. This character makes this species relatively straightforward to diagnose from other *Leptopilina*.

### Biology: Virulence assessment

To test the virulence of *Leptopilina n. sp.* on *Drosophila* host species, infection vials were established at a wasp-to-larvae ratio of 1:10 (typically 5 female wasps and 50 drosophila larvae). Two isofemale lines from two *Drosophila* species that were collected from the island of Saint Lucia were assessed: *Drosophila willistoni* (line STL3 iso3) and *Drosophila antillea* (line STL3 iso1). To collect the age-matched larvae, 20-30 male and female adult flies were placed in embryo collection cages. The embryo collection plates (60 x 15mm) contained 5mL of *Drosophila* banana food medium (Archon Scientific) and a small amount of yeast paste (active dry yeast mixed with water). The embryo collection cages were maintained in an incubator set at 25°C and 75% humidity on a 12h/12h light/dark cycle. Embryos from a restricted 12-hour egg-laying window were allowed to develop in the incubator for 3 days until they reached the size of 2^nd^ instar larvae. Larvae were carefully gently collected using extreme caution so as to not damage them.

Fifty larvae were placed into individual vials containing *Drosophila* banana food and a small cotton dental plug that was inserted into the center of the food. The cotton plug was necessary to provide a place for pupariation for the *D. antillea* species (which avoids pupariation on the side of the culture vials). Five naïve *Leptopilina n. sp.* female wasps were placed into each infection vial and were allowed to infect the larvae for 24 hours. During the infection period, the vials were closed with Flugs (Genesee Scientific cat.no.49-102) dampened with honey and water (1:1 mixture) to ensure that the wasps had access to food. After the infection period, the wasps were removed, and the vials were incubated as described above. Uninfected mock control vials were treated in the same way but were not exposed to wasps. The vials were assessed daily for the emergence of adult insects. Adult flies eclosed after 10-14 days, and adult wasps eclosed after 25-30 days. Any newly eclosed adult flies and wasps were counted and removed from the vials. The difference between the total number of adults and the initial number of larvae in each vial accounts for larval mortality.

### Biology: Western blots for *Wolbachia* detection

Active *Wolbachia* infection was assessed with western blot following published protocols [18]. Positive and negative controls included individual *Drosophila melanogaster* from previously validated *Wolbachia*-infected and –uninfected stocks (RRID: BDSC_145) [19]. Individual insects were homogenized with a pestle in 1.7 ml microcentrifuge tubes containing lysis buffer (150mM NaCl, 1% Triton X-100, 50mM TrisHCl (pH8) containing HALT protease inhibitor cocktail (Thermo Scientific) and 5 mM EDTA).

Homogenates were pelleted via centrifugation (8,000Xg for 1 minute) and the supernatant was incubated for 5 minutes at 95°C in Laemmli sample buffer with 5% β-mercaptoethanol (Bio-Rad). Proteins were separated via SDS-PAGE on 4–20% Tris-Glycine gels (NuSep) in 1X Tris/Glycine/SDS running buffer (Bio-Rad) alongside a PageRuler prestained protein ladder (Thermo Scientific), transferred to a PVDF membrane in Tris-Glycine transfer buffer with 15% methanol for 3 hours at 40v, and blocked for 5 minutes in Starting Block T20 (Thermo Scientific). The membrane was co-incubated in primary antibodies overnight at 4°C. After washing, the blot was incubated with the secondary antibody anti-mouse:HRP (Invitrogen) at 1:5000. Finally, HRP was detected with SuperSignal West Pico Chemiluminescent Substrate (Thermo Scientific). Primary antibodies included (1) mouse monoclonal Anti-Wolbachia Surface Protein (WSP), NR-31029 from BEI Resources, NIAID, NIH at 1:10,000, and (2) mouse anti-actin monoclonal at 1:20,000 (Seven Hills Bioreagents) as a loading control.

### Biology: Determining reproductive mode

To test whether the observed parthenogenesis was induced by infection with a bacterial symbiont, *Leptopilina n. sp.* female wasps were fed with the antibiotic rifampicin mixed into honey (100 mg of rifampicin powder (Sigma-Aldrich cat.no. R3501-250MG) with 1 mL of honey) and reproductive biology was assessed following standard protocols [20]. To feed the wasps, a Flug was dipped into water to dampen one side, onto which a small amount of the rifampicin-honey mixture was spread (approx. 10-20 μL). A fresh Flug with rifampicin-honey was replaced once a day for six days. To prepare vials of fly larvae, adult *Drosophila* were allowed to seed vials for 5 days, after which the adults were removed. After the six days of rifampicin treatment, the treated wasps were placed into the vials with fly larvae and allowed to infect as many larvae as possible. Wasp offspring eclosed after 25-30 days and all female and male offspring were collected and counted. Rifampicin treatment trials were performed with larval hosts from fly species that were collected on the island of Saint Lucia, either *D. melanogaster* or *D. willistoni*. Regardless of the host species, the sex ratio of the wasp offspring in each generation was always almost all female or almost all male (see results, Figure 5), and therefore the data from the trials with different host species were combined. The treatment conditions of each generation were as follows: G_1_ unmated females received 6 days of rifampicin treatment. The G_2_ offspring were almost all female and also received 6 days of rifampicin treatment, after which they produced F_0_ offspring. In order to cross *Wolbachia* depleted males and females together, a separate generation of G_1_ females was treated with rifampicin to produce G_2_ females that would eclose at the same time as the F_0_ males. These G_2_ females then received 6 days of rifampicin treatment and were then mated to F_0_ males. Subsequent generations (F_1_ – F_5_) were sexed and propagated by attempting sibling mating (when possible, see results) without further rifampicin treatments.

### Genomics: Oxford Nanopore library preparation and sequencing

DNA was extracted from a pool of ∼100 adult female wasps using a Gentra PureGene Tissue Kit (Qiagen). 3 ug of DNA was end prepped using NEBNext FFPE Repair Mix (M6630), NEBNext Ultra II End repair/dA-tailing Module (E7546), and NEBNext Quick Ligation Module (E6056) and subsequently library prepped using the ONT supplied SQK-LSK110 kit. Sequencing took place on an Oxford Nanopore Minion over 3 days with nuclease wash followed by reloading after 24 hours. The sequencing computer was built with an AMD Ryzen 3900x processor with 12 threads, 24 cores, 64 Gb RAM and a 1 Tb SSD, and a GeForce 2080Ti with 4352 CUDA cores running Ubuntu Linux 20.04. We monitored the CUDA cores with nvtop (https://github.com/Syllo/nvtop). Fastq files were generated from the Oxford Nanopore Minion instrument and called with Guppy v6.1.5. We used the GPU enabled version of guppy in concert to enable live basecalling with the fast base calling model and for post-hoc base calling with the super (sup) accuracy model called via Minknow v22.05.5. All resulting fastq.gz files were concatenated and used for further processing.

### Genomics: Illumina sequencing

Total DNA was extracted from a pool of ten adult wasps using the DNeasy blood and tissue kit (Qiagen). Sequencing library preparation was performed with the NEBNext Ultra II DNA library preparation kit (New England Biolabs) according to the manufacturer’s protocols including 10 cycles of PCR amplification. The resulting library was subjected to 150-bp paired-end sequencing on the Illumina NextSeq 500 platform at the Indiana University Center for Genomics and Bioinformatics (Bloomington, IN) to generate 181,561,108 read pairs. To aid in gene annotation we also sequenced wasp transcriptomes. Total RNA was extracted from a flash-frozen, mixed stage pool of 10 wasps (adults, pupae, larvae) using the Monarch® Total RNA Miniprep Kit (New England Biolabs) including the DNase treatment step. An RNA-Seq library was prepared with the Illumina TruSeq Stranded mRNA Library Prep Kit, including rRNA removal with oligo-dT beads, and sequenced on a NovaSeq with S4 chemistry at the University of Minnesota Genomics Center to generate 64,677,726 150-bp paired-end reads.

### Genomics: Nuclear genome assembly, curation, and quality control

All reads greater than 5kb were filtered with SeqKit v2.5.1 [21], assembled with Flye v2.9 [22], and the assembly was polished with Medaka v1.5.0 (https://github.com/nanoporetech/medaka). Haplotigs were purged with PurgeHaplotigs v1.1.2 [23]. Pilon v.122 [23] was used for polishing: first WGS Illumina reads were mapped to the long-read assembly with bwa v0.7.17 [24], alignment files were converted and sorted with samtools v1.10 [25], and finally pilon was called for polishing to generate an updated assembly. Four rounds of pilon polishing were performed, each time mapping the short reads to the most recent pilon output prior to pilon correction. Manual curation of the assembly and identification of cytoplasmic genomes was informed by results derived from running blobtools [26]. We discarded contigs that were deemed anomalous for the following reasons: (1) less than 5,000 bp in length with no BUSCOs (because >5k reads were used during assembly, these are very likely to be chimeric and/or haplotigs), or, (2) having blast hits to non-arthropod taxa across the length of the contig, unusual sequencing coverage(<15X; >800X), and unusual GC content (outside of 23-33%). Finally, we checked the previously purged contigs and recovered those that appeared to encode for *Leptopilina* rRNAs. Assemblies were assessed with BUSCO v.5.3.2 [27] against the hymenoptera_odb10 database in genome mode (‘-m genome’).

### Genomics: Methylation

Nanopore runs were base called with guppy v6.3.8 using model “dna_r9.4.1_450bps_modbases_5mc_cg_sup” to call 5-methylcytosines in a CpG context only. Aligned bam reads containing modified base information were converted into bed format with modbam2bed, with strict parameters on canonical and modified base probabilities where bases are only called as canonical or modified if they have a greater than 95% probability estimate. Parameters: modbam2bed –m 5mC –t 32 –e –p total.modmapped –-aggregate –-cpg dtwasp-final-curated.fa total.modmapped.bam –a 0.05 –b 0.95 > total.modmapped.bed”. Global methylation was calculated by an awk script.

### Genomics: Mitogenome

Mitogenome annotation with MITOS2 [28] and manual rearrangement to start the circular mitogenome at Cox1 (as is convention) was done as in [29]. MITOS2 parameters included the RefSeq 63 Metazoa reference and the invertebrate mitochondrial translation code. Manual curation was facilitated by comparison to published mitochondrial genomes from *Leptopilina boulardi* (KU665622.1) and *Leptopilina syphax* (MT649407.1). A phylogeny of *Leptopilina* was constructed based on COX1 (“COI”) sequences available from the DROP database [3]. A subset of the available *Leptopilina* sequences were selected based on (1) sampling across the genus, (2) limiting the total number of sequences for any given species, (3) length of the barcoding region sequenced, and (4) checking for open reading frame. In total 55 sequences were used for phylogenetic reconstruction. These include an outgroup (*Ganaspis kimorum*), previously sequenced barcodes for the *Leptopilina n. sp.* STL6 colony [3], and the COI sequence derived from the mitogenome assembly. Nucleotide sequences were aligned with MAFFT v7.511 [30] using the L-INS-i alignment algorithm and a five point gap open penalty (-op 5). Comparison to the translated amino acid alignment verified that codon relationships were maintained and we did not need to back-translate an amino acid alignment. Ends of the nucleotide alignment were trimmed such that all sequences were represented at all positions. Phylogenetic reconstruction was then performed with IQtree v1.6.11 [31] using a pre-analysis model optimization step to select the final model (here, TIM3+F+I+G4) plus 1000 “ultrafast” bootstraps.

### Genomics: Repeat identification

Repetitive DNA content was estimated using RepeatModeler2 [32] using the curated Dfam database release 3.5 July 2021 [33], combined with the Repbase RepeatMasker database downloaded on October 26, 2018. RepeatMasker was run with the newly identified library to annotate the assembly with the following parameters: RepeatMasker v4.1.2-pl –pa 24 –s [sensitive mode] [using rmblast 2.11.0+] –lib ultimatefamilies.fa ultimateGenome.fasta. Each repeat library was also mapped against all other species genomes with the following parameters: RepeatMasker v4.1.2-pl –pa 24 –s [sensitive mode] [using rmblast 2.11.0+] –lib families.fa genome.fasta.

### Genomics: Gene annotation and ortholog clustering

Gene models were predicted with GeMoMa v1.9 [34] and leveraged three lines of evidence: (1) RefSeq annotations for *Leptopilina heterotoma* (GCF_015476425.1) (2) RefSeq annotations for *Leptopilina boulardi* (GCF_019393585.1) and (3) RNA-seq reads derived from the mixed pool of wasps. RNA-seq reads were mapped to a hard masked version of the reference genome using STAR v2.5.3a [35]. GeMoMa was called with the following additional parameters: tblastn=TRUE, AnnotationFinalizer.n=false, AnnotationFinalizer.r=SIMPLE, AnnotationFinalizer.t=true, AnnotationFinalizer.p=LMAL, pc=true, o=true. To generate annotations for *Leptopilina clavipes* and *Ganaspis kimorum*, we ran the same annotation pipeline on hard-masked assemblies, only without the RNA-seq data. Protein sequences from each genome were clustered into orthologous groups using OrthoFinder v.2.5.4 [36]. For genes represented by multiple transcript variants, only the longest isoform was used in clustering.

### Genomics: *Wolbachia* genome

A single contig was identified to be a *Wolbachia* genome, based on size, circularity, and Blobtools results. Genome completeness was assessed with BUSCO v5.3.2 [27] against the rickettsiales_odb10 database in genome mode (‘-m genome’). Genome annotation and phylogenetics were performed by running the *Wolbachia* Phylogeny Pipeline (WHOP; https://github.com/gerthmicha/WHOP [37]). In brief, the pipeline takes a provided database of bacterial genomes (*Wolbachia* strains and Rickettsiales outgroups) plus additional genomes provided by the user, annotates them with Prokka v1.14.6 [38], clusters protein sequences into orthogroups with OrthoFinder v2.5.4 [36], aligns the single-copy orthologs with MAFFT L-INS-i v7.487 [30], removes recombining genes with PhiPack v1.1 [39], and then concatenates the alignments prior to phylogenetic reconstruction with IQtree v2.2.3 [31]. IQtree was first run with a pre-analysis model optimization step (-m LG+I+G4 –pre –redo –fast), followed by final reconstruction with the selected model and 1000 bootstrap replicates. The *w*Lmal genome was re-oriented to start at *dnaA* and proksee (https://proksee.ca/ [40]) was used for visualization. Prophage regions were identified with VirSorter2 v.2.2.4 [41], and Phigaro v.2.3.0 [42], and other mobile elements were identified with mobileOG-db v.1.0.1 [43], all implemented in proksee with default parameters. To aid in manual annotation of *w*Lmal we leveraged the orthogroup relationships between *w*Lmal and *w*Mel generated by WHOP to check for specific proteins. Additionally, we used blastp and tblastn with default parameters to query against the protein sequences or genome, respectively, for a select number of functionally relevant *Wolbachia* proteins not found in *w*Mel. All manual annotation queries and matches are in Supplemental Tables S5-6.

### Data visualization

Data visualization was carried out in R version 4.1.3 [44] and leveraged Paul Tol’s “Medium-contrast qualitative colour scheme”, (R: khroma [45]). Newick files from phylogenetic reconstructions were visualized in FigTree v1.4.4 (https://github.com/rambaut/figtree/) and annotated in Inkscape. Proksee (https://proksee.ca/ [40]) was used for genome visualization, and the r package ‘gggenes’ v.0.5.1 was used to plot gene models.

## RESULTS

### Collections on Saint Lucia

Banana and calabasa fruit traps were used to collect *Drosophila* and their associates on Saint Lucia in 2019. The calabasa traps at STL3 yielded six female *Drosophila antillea* and one female *Drosophila willistoni.* Isofemale lines of *D. antillea* and *D. willistoni* were established from single females that originated from the STL3 collection site. At site STL6, banana traps contained a total of 12 *melanogaster*-like flies and 1 *repleta* group fly. Calabasa traps contained 4 *melanogaster*-like flies, 4 *repleta* group flies and 20 *dunni*-group flies. Wasp “STL6” was collected from a calabasa trap on the morning of 1 March 2019: a clear day with a high temperature of 27C (Figure 1). The STL6 wasp was placed in a vial with developing *Drosophila willistoni* (from the STL3 site, see above) to initiate a culture, and then shipped back to Indiana University where they have been maintained since (see methods).

### *Leptopilina n. sp.* Buffington, Lue, Davis & Tracey sp. nov

#### Species Diagnosis

*Leptopilina n. sp.* is distinguishable from other *Leptopilina* species by a number of characters, including: flagellomere colors, in which F1-F4 are honey-brown, F5-F10 are dark brown, and F11 is yellow (Figure 2A); the typical flagellomere color in *Leptopilina* do not match the unique combination here. Latero-posterior margin of pronotal plate is with elongate, golden setae that reach the anterior margin of the mesoscutum (Figure 2B (LM) and Figure 3A-B (SEM)). No other known *Leptopilina* have setae so long off the pronotal plate (Figure 2B); other species have hairs here, but not so long. Members of the *clavipes* group *sensu* [15] tend to also have sparse, scattered setae over the mesoscutum, whereas *Leptopilina n. sp.* is entirely glabrous (Figure 3A-B).

**Figure 2.**
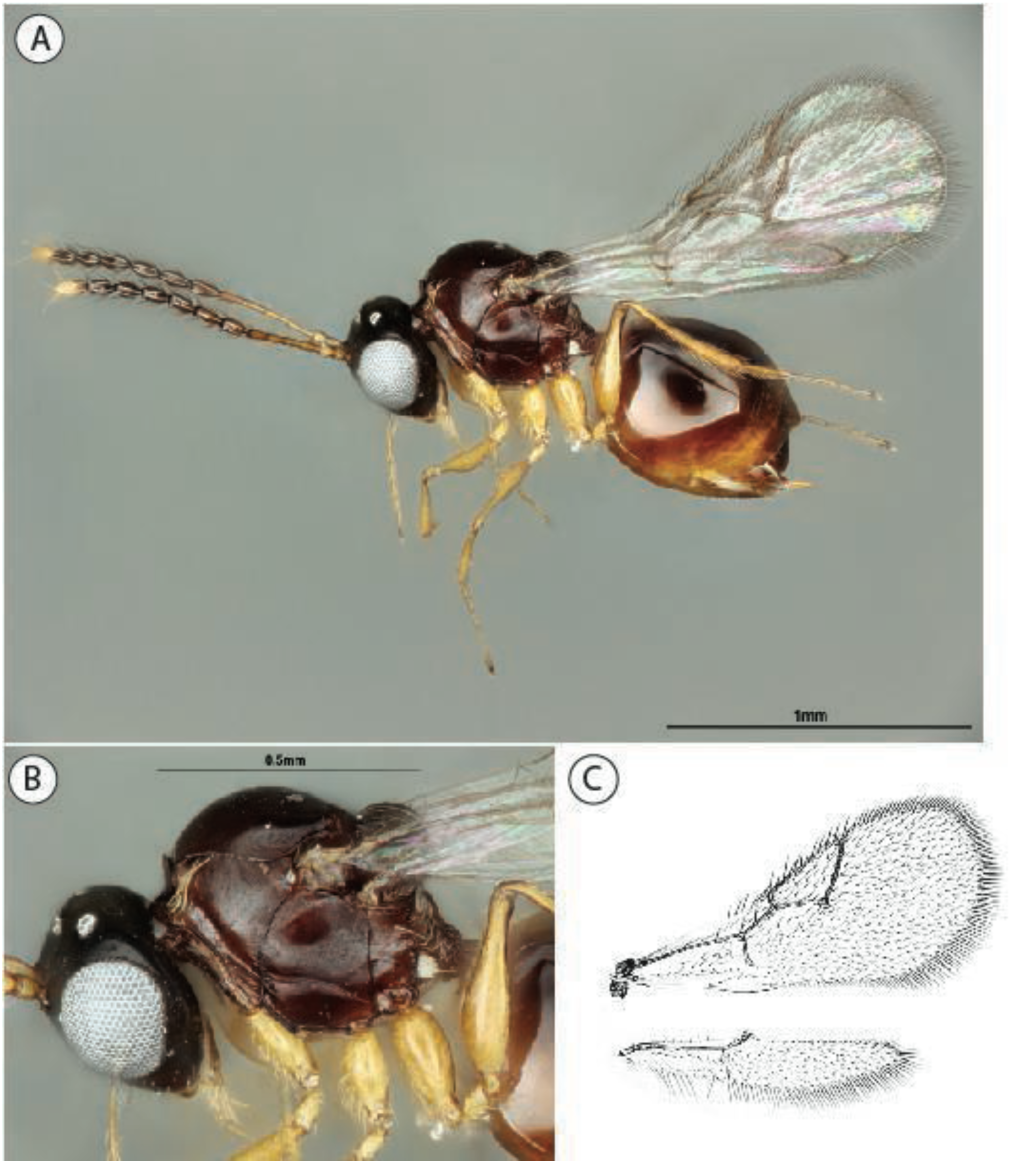
Leptopilina n. sp. **(A)** Lateral view, **(B)** lateral view of mesosoma, and **(C)** wing vein pattern.

**Figure 3.**
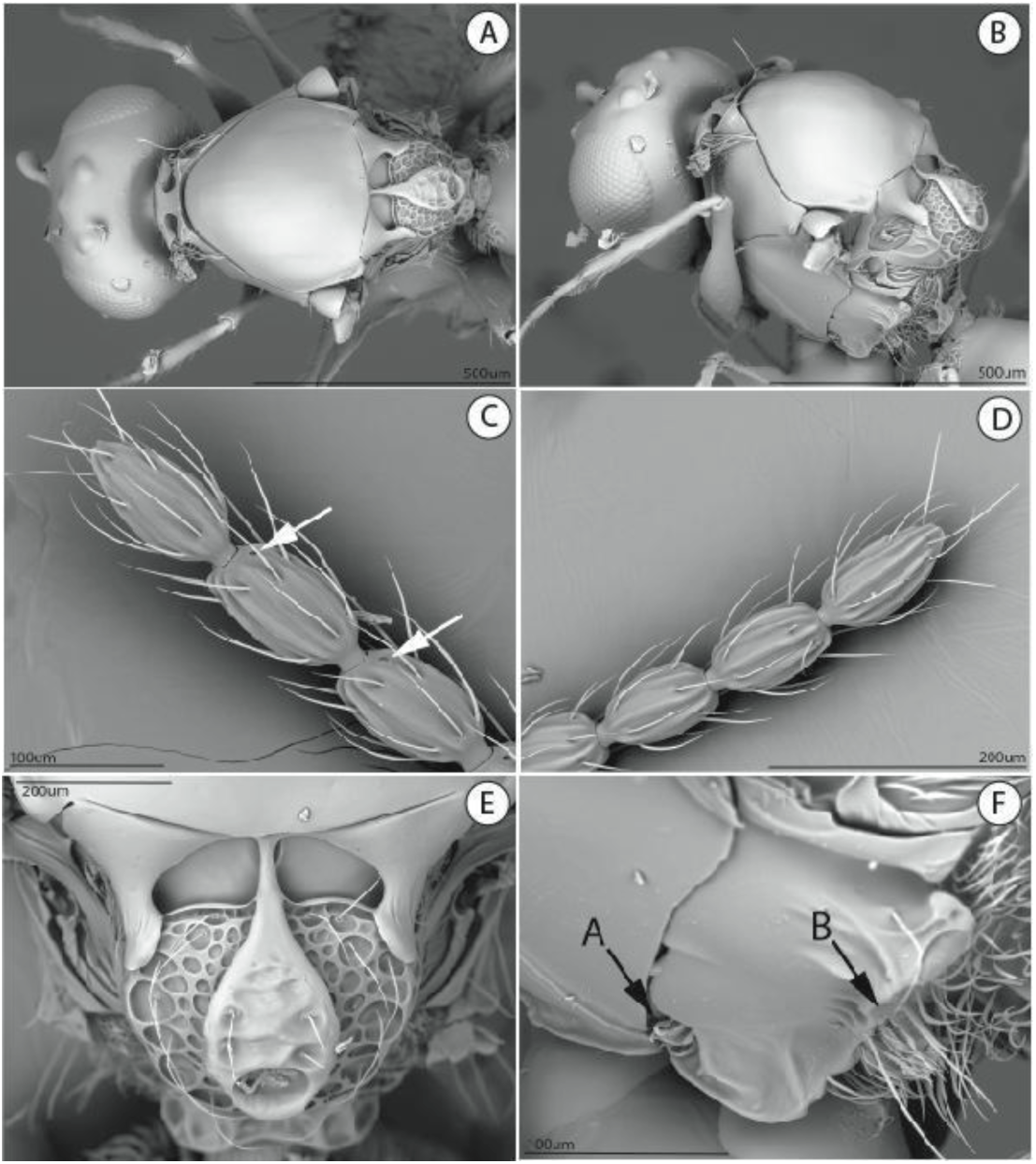
SEM images of *Leptopilina n. sp.* **(A)** Top view of mesoscutum, **(B)** Lateral-posterior view of mesosoma, **(C)** campaniform sensillae, white arrow, compared to **(D)** no campaniform sensillae, **(E)** scutellum plate, and **(F)** metepimeron.

### Morphological Description

Coloration with head, mesosoma, metasoma black to dark brown, legs light brown. Sculpture on vertex, lateral surface of pronotum and mesoscutum absent, surface smooth. *Antenna*. Female antenna composed of scape, pecidel, 11 flagellomeres (F1-F11). F1-F4 sub-cylinderical, slightly expanded distally, smooth, non-claval segments. F5-F11 claval segments, abruptly broader than non-claval segments, distinctly barrel-shaped, with slightly more erect, dense setae; claval segments with placoidal sensillae distributed evenly around each segment. Single campaniform sensillae present ventro-apically on F6-F10 (arrow, Figure 3C; compare to Figure 3D). Articulation between flagellomeres in antenna moniliform, segments distinctly separated by narrow neck-like articulation. Male antenna composed of 13 flagellomeres, F1 as long as F2, slightly curved and excavated distally; campaniform sensillae on all flagellomeres.

*Head*. In anterior view, broadly triangular. Pubescence on head sparse, single setae scattered over face. Sculpture along lateral margin of occiput absent. Gena (measured from compound eye to posterolateral margin of head) short, ratio of length of gena to length of compound eye in dorsal view <0.3. Sculpture of gena absent, smooth. Lateral margin of occiput evenly rounded, not well defined. Occiput (except extreme lateral margin) smooth. Carina issuing from lateral margin of postocciput absent. Ocelli small, ratio of maximum diameter of a lateral ocellus to shortest distance between lateral ocelli 0.2-0.4. Anterior ocellus far from posterior ocelli, clearly separate anterior ocellus to anterior margins of posterior ocelli. Relative position of antennal sockets intermediate, ratio of vertical distance between inner margin of antennal foramen and ventral margin of clypeus to vertical distance between anterior ocellus and antennal rim 2.0-4.0. Median keel absent. Vertical carina adjacent to ventral margin of antennal socket absent; facial sculpture absent, surface smooth. Facial impression absent, face flat. Antennal scrobe absent. Anterior tentorial pits small. Vertical delineations on lower face absent. Ventral clypeal margin laterally, close to anterior mandibular articulation, straight. Ventral clypeal margin medially straight, flat, with 4-6 setae overhanging base of mandibles. Clypeus smooth. Malar space adjacent to anterior articulation of mandible evenly rounded, smooth. Malar sulcus present, composed of single groove. Eye removed from lateral ocelli, ratio of distance between compound eye and posterior mandibular articulation to distance between posterior ocellus and compound eye <1.2. Compound eyes, in dorsal view, moderately protruding from the surface of the head. Pubescence on compound eyes absent. Orbital furrows absent. Lateral frontal carina of face absent. Dorsal, posterior aspect of vertex smooth. Hair punctures on lateral aspect of vertex absent. Posterior surface of head deeply impressed around postocciput.

*Mouthparts*. Apical segment of maxillary palp with pubescence, consisting of one long erect setae. Apical seta on apical segment of maxillary palp longer than twice length of second longest apical seta. Maxillary palp composed of four segments. Last two segments of maxillary palp (in normal repose) straight. Apical segment of maxillary palp about 1.5 times as long as preceding segment. Left mandible with three teeth; scattered setae dorsally.

*Mesosoma.* Long auburn-golden setae present, originating from anterior inflection of pronotum, extending to anterior margin of mesoscutum; rest of pronotum glabrous (Figure 3A-B). Anteroventral inflection of pronotum narrow. Pubescence on lateral surface of pronotum absent. No ridges present posterior to pronotal plate. Anterior flange of pronotal plate distinctly protruding anteriorly, completely smooth. Ridges extending posteriorly from lateral margin of pronotal plate absent. Lateral pronotal carina absent. Crest of pronotal plate absent. Dorsal margin of pronotal plate (in anterior view) spatulate. Submedian pronotal depressions open laterally, deep. Lateral margin of pronotal plate defined all the way to the dorsal margin of the pronotum. Width of pronotal plate narrow, not nearly as wide as head.

Mesoscutal surface convex, evenly curved. Sculpture on mesoscutum absent, entire surface smooth, shiny, with sparse long hairs. Notauli absent. Median mesoscutal carina absent. Anterior admedial lines absent. Median mesoscutal impression absent. Parascutal carina nearly straight.

Mesopleuron entirely smooth. Subpleuron entirely smooth, glabrous. Lower pleuron entirely smooth, glabrous. Epicnemial carina absent. Lateroventral mesopleural carina present, marking abrupt change of slope of mesopectus. Mesopleural triangle absent. Subalar pit not visible. Speculum absent. Mesopleural carina present, complete, composed of one complete, uninterrupted carina. Anterior end of mesopleural carina inserting above notch in anterior margin of mesopleuron; posterior end terminates at setal pit of lower metapleuron.

Dorsal surface of scutellum foveate-areolet (Figure 3E). Circumscutellar carina present, complete, delimiting dorsal and ventral halves of scutellum. Posterior margin of axillula marked by distinct ledge, axillula distinctly impressed adjacent to ledge. Latero-ventral margin of scutellum, posterior to axillula, weakly rugulose. Dorsoposterior part of scutellum rounded. Transverse median carina on scutellar plate absent. Dorsal part of scutellum entirely areolate. Scutellar plate, in dorsal view, medium sized, exposing about half of scutellum, gently foveate with three to four single setae; glandular release pit at posterior end, oval. Scutellar fovea present, two, distinctly margined posteriorly, smooth. Longitudinal scutellar carinae absent. Single longitudinal carina separating scutellar foveae present, elongage, ending past posterior margin of foveae, gently expanding into teardrop-shaped scutellar plate. Postero-lateral margin of scutellum rounded. Lateral bar smooth, narrow.

Posterior impression of metepimeron absent. Metapectal cavity anterodorsal to metacoxal base present, well-defined (Figure 3F, arrow A). Anterior margin of metapectal-propodeal complex meeting mesopleuron at same level at point corresponding to anterior end of metapleural carina. Posteroventral corner of metapleuron (in lateral view) not extended posteriorly. Anterior impression of metepimeron absent. Posterior margin of metepimeron distinct, separating metepimeron from propodeum, with distinct ridge (Figure 3F, arrow B). Anteroventral setal pit of lower metapleuron present, with foamy setae.

Subalar area broadened anteriorly, narrowed posteriorly. Prespiracular process present, blunt, lobe-like, polished, at end of polished prespiracular groove; posteroventral corner of prespiracular groove extended to distinct lobe. Dorsellum absent. Anterior impression of metepisternum, immediately beneath anterior end of metapleural carina, present. Pubescence consisting of few hairs on posterior part of metepisternum, few sparse hairs on propodeum, but with distinct posterior swatch of foamy setae at midline of metapleuron.

Pubescence posterolaterally on metacoxae sparse, with a single postero-dorsal button of dense pubescence. Microsculpture on hind coxa absent. Longitudinal ridge on the posterior surface of metatibia absent. Metafemoral tooth present, elongate, with adjacent serrate ridge posteriorly. Ratio of first metatarsal segment to remaining 4 segments less than 10; metatarsus 5 distinctly elongate relative to 2 through 4.

Wing vein M absent. Pubescence of fore wing present, long, moderately dense on most of surface. Apical margin of female fore wing rounded. Rs+M of forewing defined but nebulous at point of origin from basal vein at posterior third. Mesal end of Rs+M vein situated closer to anterior margin of wing, directed towards middle of basalis. Vein R1 forming marginal cell completely. Basal abscissa of R1 (the abscissa between 2r and the wing margin) of fore wing as broad as adjacent wing veins. Coloration of wing absent, entire wing hyaline. Marginal cell of fore wing membranous, similar to other wing cells. Areolet absent. Hair fringe along apical margin of fore wing present, long or very long (Figure 2C).

Propodeal spurs absent. Lateral propodeal carinae present, not reaching scutellum. Ventral end of lateral propodeal carina reaching nucha, carinae separated from each other, elbowed at midpoint, not parallel. Inter propodeal carinae space lightly setose, underlying surface appearing smooth. Petiolar rim of uniform width along entire circumference. Petiolar foramen removed from metacoxae, directed posteriorly. Horizontal carina running anteriorly from lateral propodeal carina absent, moderately setose. Calyptra, in lateral view, rounded. Propodeum not strongly drawn out posteriorly. Calyptra, in posterior view, dorsoventrally elongate or rounded.

*Metasoma.* Petiole about as long as wide. Surface of petiole longitudinally costate, ventral keel absent. Posterior part of female petiole abruptly widened. Ventral and lateral parts of petiolar rim broad.

Setal band (hairy ring) at base of tergum 3 present, interrupted dorsally, ventrally. Tergum 3 indistinct, fused with syntergum. Posterior margin of tergum 3 indistinct, fused with tergum 4 in syntergum. Posterior margin of tergum 4 evenly rounded. Sternum 3 encompassed by syntergum. Sculpture on metasomal terga absent. Syntergum present with terga 3 to 5 fused, ventral margin rounded. Peglike setae on T6-T7 absent. Postero-ventral cavities of female metasoma T7 present, glabrous save for few, long setae. Female postero-ventral margin of T6-T7 straight, parallel. Terebrum and hypopygium (in lateral view) curved, pointing upward. Ovipositor clip present.

#### Etymology

The species epithet refers to locations on Saint Lucia: Malgretoute, a town near the island’s active volcano, and the near-by Malgretoute Beach, from which the iconic Saint Lucia Pitons are visible.

#### Material examined. Holotype

Female. Saint Lucia, Barr d’Isle trail, central. lat 13.924, long –60.959, elevation 308.7m. 17.II.2019, Jeremy S. Davis, collected from calabasa bait trap. USNMENT01025555. [1] Actual specimen taken from Indiana University colony derived from the above collection locality.

#### Paratypes

9 females, 3 males. Same collection data as holotype. USNMENT01557161, USNMENT01025556, USNMENT01025557, USNMENT01025554, USNMENT01557162, USNMENT01557158, USNMENT01448469, USNMENT01448467, USNMENT01932975-USNMENT01932978.

#### Other material examined

8 females. INRA laboratory, Petit Bourg, Guadeloupe, January 1982. Carton, coll. USNMENT01025775-USNMENT01025782

### *Leptopilina n. sp*. is highly virulent on *D. antillea* flies

As noted extensively above, our laboratory cultures of *Leptopilina n. sp.* were collected on Saint Lucia. This island is also the only reported location for the *Drosophila antillea* species of the dunni subgroup. To our knowledge, the susceptibility of dunni group *Drosophila* to parasitoid wasps has not been previously investigated. Thus, we wondered if the *D. antillea* flies would be susceptible, or if they were possibly resistant, to infection by a wasp species that is also found on Saint Lucia. We allowed female wasps to oviposit for 24 hours on larval cultures that were derived from isofemale lines of either *D. antillea* or *D. willistoni* (that both originated from the STL3 collection site) and then determined the proportion of wasps and flies that emerged from the infection vials. The results clearly indicated a high degree of virulence of *Leptopilina n. sp.* on *D. antillea* (76.5% ± 5.6% wasp eclosion) relative to *D. willistoni* (20% ± 3.8% wasp eclosion) as host species (Figure 4).

**Figure 4.**
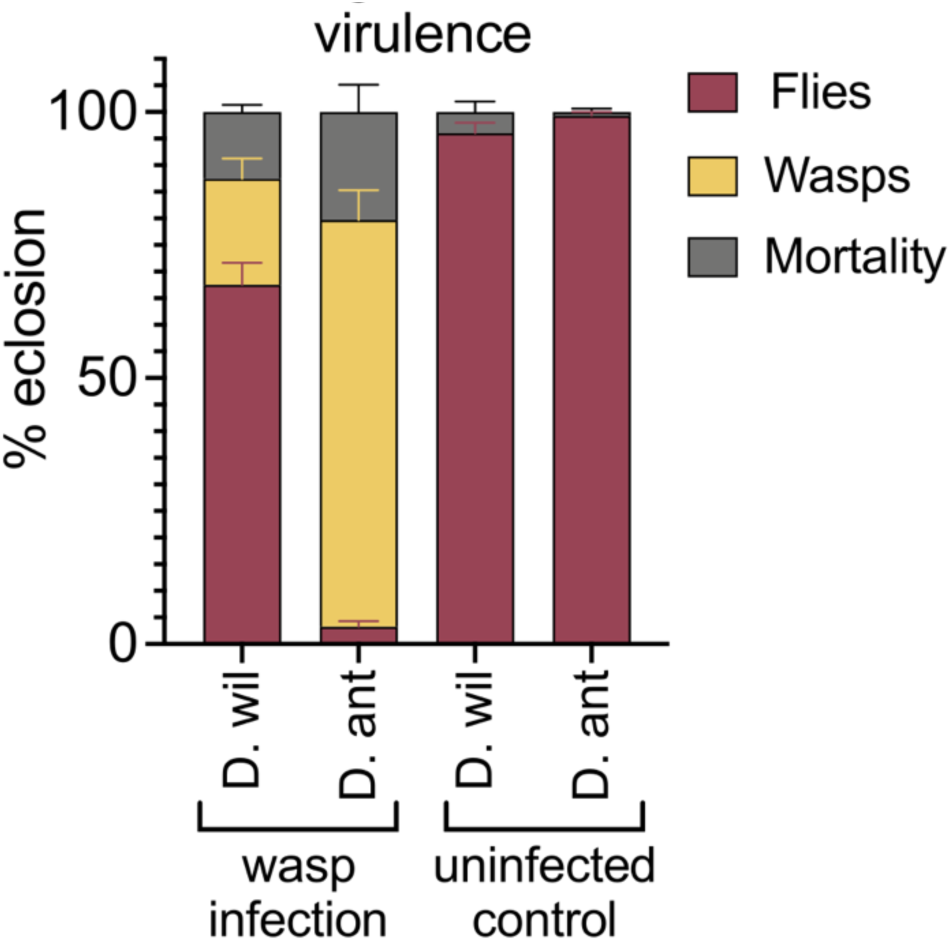
*Leptopilina n. sp.* shows high virulence on a host fly species that is exclusive to Saint Lucia. Percentage of adult flies and wasps that eclosed from vials infected by *Leptopilina n. sp.,* and from uninfected control vials. Mortality represents the percentage of larvae that did not develop to adulthood. *L. mal.* is clearly more virulent on *Drosophila antillea* (“D. ant”) hosts than on a *Drosophila willistoni* (“D. wil”) host strain that was also collected on Saint Lucia. Data are plotted as mean ± SEM; n = 8 infection vials, n = 3 uninfected control vials.

**Figure 5.**
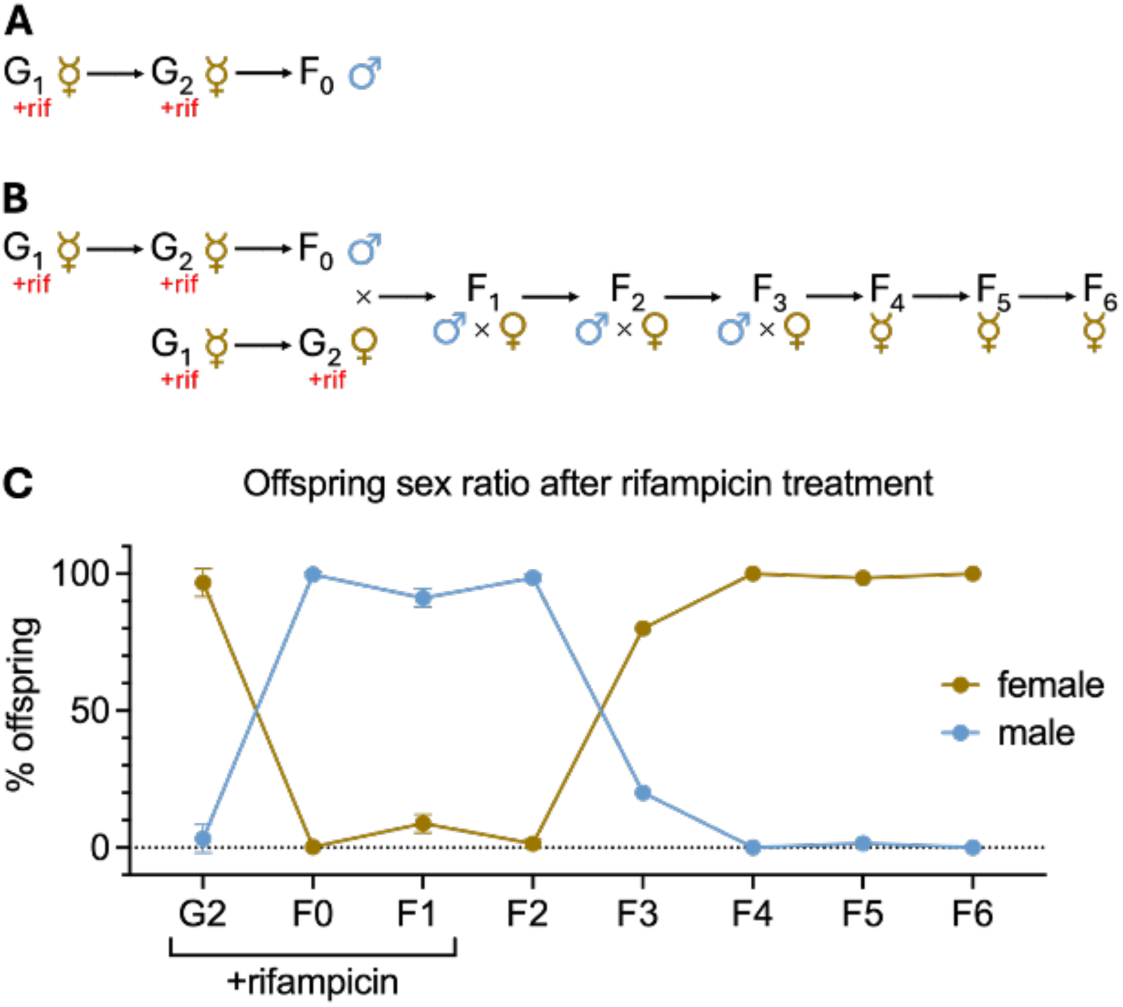
*Leptopilina n. sp.* reproduces by symbiont-mediated thelytokous parthenogenesis. **(A)** Schematic depicting the production of male progeny after two generations of antibiotic treatment (“+rif”), which supports the hypothesis of symbiont mediated parthenogenesis. **(B)** Schematic of antibiotic treatment and crossing scheme to test for sexual reproduction after symbiont depletion, and an attempt to generate a sexual, symbiont free strain. **(C)** Percentage of female and male offspring in each generation. Data is plotted as mean ± SEM. For generations G_2_ to F_2_, n = 4-8 infection vials; for generations F_3_ to F_6_, n = 1 infection vial.

### *Wolbachia* mediates obligate thelytokous parthenogenesis in *Leptopilina n. sp*

While maintaining *Leptopilina n. sp.* wasp lines we noticed that the cultures were primarily female (>95%). And, in one generation of passaging, the cultures became entirely female. Fearing that we would lose the culture, we infected flies with these female wasps in an attempt to obtain haploid male offspring that could be mated back to their mothers to obtain more diploid females. However, the cultures generated by the virgin female wasps were again entirely female. This indicated that this species of wasp was capable of asexually producing females (*i.e.*, thelytokous parthenogenesis). Furthermore, the observation that males were rarely obtained in the cultures was unlikely to be strictly a consequence of the wasps having especially high rates of fertilization (which gives rise to female offspring in arrhenotokous, sexual, haplodiploids). In many parasitic wasp species, including a suite of other *Leptopilina sp.*, thelytokous parthenogenesis is mediated by vertically transmitted bacterial symbionts such as *Wolbachia* (α-proteobacteria: Rickettsiales) [46–48]. We tested for the presence of *Wolbachia* infection by Western blot, targeting a *Wolbachia* specific protein and found that indeed the wasps were infected with this microbe (Supplemental Figure S1).

To determine if a microbe was responsible for the asexual production of females, we used antibiotic treatment with rifampicin to remove the infection and assess changes in the offspring sex ratio derived from unmated females (Figure 5A). After treating unmated adult females (G_1_) with rifampicin, they were allowed to parasitize host larvae. Their broods (G_2_) were almost entirely female (98.6% ± 2.6% female). Critically, *Leptopilina* are largely proovigenic: all their eggs develop and mature during the pupal period [49]. This means that symbiont-mediated effects are likely established much earlier, and treating adults is “too late” (as has been seen in other wasps [50]). Thus, we performed a second generation of rifampicin treatment: G_2_ rifampicin-treated females produced offspring (F_0_) that were almost all male (99.7% ± 0.3% male), a clear indicator of symbiont-mediated parthenogenesis.

We attempted to generate a symbiont-free sexually reproducing line of wasps (Figure 5B) but were unsuccessful. Specifically, symbiont depleted males and treated G_2_ females were crossed to assess sexual reproduction, but offspring continued to be almost all male (F_1_-F_2_, 91.2% ± 1.5% and 98.5% ± 0.6% male, respectively), indicating no sexual reproduction (Figure 5C). After antibiotic treatment was discontinued, rare female progeny that were allowed to reproduce produced sex ratios that gradually returned to being almost entirely female (F_3_-F_4_, 80% and 100% female, respectively), and unmated females were again able to produce female offspring by parthenogenesis (F_5_-F_6_, 98.5% and 100% female, respectively).

### *Leptopilina n. sp.* genome sequencing and assembly

We generated a genome assembly for *Leptopilina n. sp.* based on nanopore long read sequencing (Table 1, Supplemental Tables S1-S3). The draft assembly with Flye leveraged all nanopore reads longer than 5,000 bp (Supplemental Table S1) and generated 1,666 contigs totaling just over 1Gbp in total length (Table 1; Supplemental Figure S2). The assembly was then polished with illumina short reads: short-read mapping rates were very high (98.9% of reads mapped). After polishing, separating cytoplasmic genomes, removing haplotigs, purging contaminants, and removing otherwise spurious contigs, the ultimate nuclear genome assembly was 1,408 contigs and ∼995Mbp in size with an N50 of >1.8Mbp (Table 1). The longest contig in the assembly was close to 13Mbp in length. Assembly quality was assessed with BUSCO. Just under 90% of the hymenopteran BUSCO proteins were complete, similar to other assemblies in the genus (Table 2). We recovered two cytoplasmic genomes: mitochondrion and *Wolbachia*. Additional low-coverage bacterial sequences that were purged from the assembly belonged to typical associates of the fly food and microbiome (Supplemental Results in File S1 and Supplemental Table S3). No other symbionts were identified in the contigs (e.g., *Cardinium*, *Spiroplasma,* etc [51–53]). Thus, we determine that *Wolbachia* is likely the cause of thelytokous parthenogenesis.

**Table 1.** Assembly statistics for *Leptopilina n. sp*.

| Metric | Draft | Ultimate |
| --- | --- | --- |
| Contigs | 1,666 | 1,408 |
| Length (bp) | 1,004,830,435 | 994,948,826 |
| Min length | 499 | 5,452 |
| Avg length | 603,134 | 706,640 |
| Max length | 12,918,385 | 12,924,377 |
| N50 | 1,831,405 | 1,835,757 |
| %GC | 29.85 | 29.75 |
| BUSCO* (Hymenoptera) | C:89.3%[S:88.8%,D:0.5%], F:2.9%,M:7.8%,n:5991 | C:89.7%[S:89.2%,D:0.5%], F:2.7%,M:7.6%,n:5991 |

\*Standard BUSCO annotation: Complete BUSCOs (C) [Complete and single-copy BUSCOs (S), Complete and duplicated BUSCOs (D)], Fragmented BUSCOs (F), Missing BUSCOs (M), Total BUSCO groups searched (n)

**Table 2.** Additional genomes used in comparative analyses.

| Species | Assembly Accession | Size (bp) | Protein Coding Genes | BUSCO* (Hymenoptera) |
| --- | --- | --- | --- | --- |
| *Leptopilina boulardi* | ZJU_Lbou_2.1 | 354,804,354 | 13,647 | C:91.4%[S:90.9%,D:0.5%], F:1.7%,M:6.9%,n:5991 |
| *Leptopilina clavipes* | ASM185565v1 | 255,343,917 | 13,920 | C:77.8%[S:77.3%,D:0.5%], F:7.3%,M:14.9%,n:5991 |
| *Leptopilina heterotoma* | ASM1547642v1 | 484,694,486 | 14,450 | C:91.1%[S:90.6%,D:0.5%], F:1.9%,M:7.0%,n:5991 |
| *Ganaspis kimorum*<sup>#</sup> | GCA_037103525.1 | 1,025,447,244 | 20,567 | C:89.4%[S:88.8%,D:0.6%], F:2.7%,M:7.9%,n:5991 |

\* Standard BUSCO annotation: Complete BUSCOs (C) [Complete and single-copy BUSCOs (S), Complete and duplicated BUSCOs (D)], Fragmented BUSCOs (F), Missing BUSCOs (M), Total BUSCO groups searched (n). Hymenoptera dataset used.
<sup>#</sup> Previously '*Ganaspis* near *brasiliensis*' (including in genome report [54]), but recently re-assigned [6].

### Methylation

We used a methyl-sensitive base-calling model to predict 5-methylcytosines in a CpG context. Across the nuclear, mitochondrial, and *Wolbachia* genomes, fractional methylation was less than 1% (Genome: 0.600%, mitochondrion: 0.547%, *Wolbachia*: 0.598%). These estimates likely do not represent low levels of true genomic methylation, rather, they are probably background noise due to error rates inherent to the sequencing and base calling.

### Mitogenome

We recovered a circular mitochondrial genome (14,490 bp; 20.6% GC; 3,562X sequencing coverage, Figure 6, Supplemental Table S3). Using the barcoding sequence region from the mitochondrial genome, plus representative sequences across the genus, we reconstructed a phylogeny of *Leptopilina sp.* (Figure 6A). *Leptopilina n. sp.* sequences were all identical, and formed a monophyletic group more than 10% divergent from the next most closely related species in the dataset, further supporting its designation as a new species. Importantly, *Leptopilina n. sp.* is sister to a clade within the *Leptopilina* genus that contains all other species for which genome sequences are available (*Leptopilina clavipes, Leptopilina boulardi,* and *Leptopilina heterotoma*). Annotation of the mitochondrial genome revealed that gene order was largely conserved as compared to other *Leptopilina* mitochondria. Additionally, we annotated the expected rRNAs and protein coding genes. However, insect mitochondrial tRNAs are often difficult to annotate [55], and in *Leptopilina n. sp.* we were unable to find tRNAs for asparagine (N), alanine (A), aspartic acid (D), pheylalanine (F), serine anticodon-1 (S1), and arginine (R). Related to that, the region between the 3’ end of the 16S rRNA (rrnL) and the 5’ end of *nad2*, was quite divergent as compared to *Leptopilina boulardi,* in which there is a cluster of tRNAs (including D, F, S1, N, A, plus glutamic acid, glycine, and histidine). It is likely that divergence and/or noncanonical tRNA structures, as is often the case in insect mitochondria, may be precluding a complete annotation.

**Figure 6.**
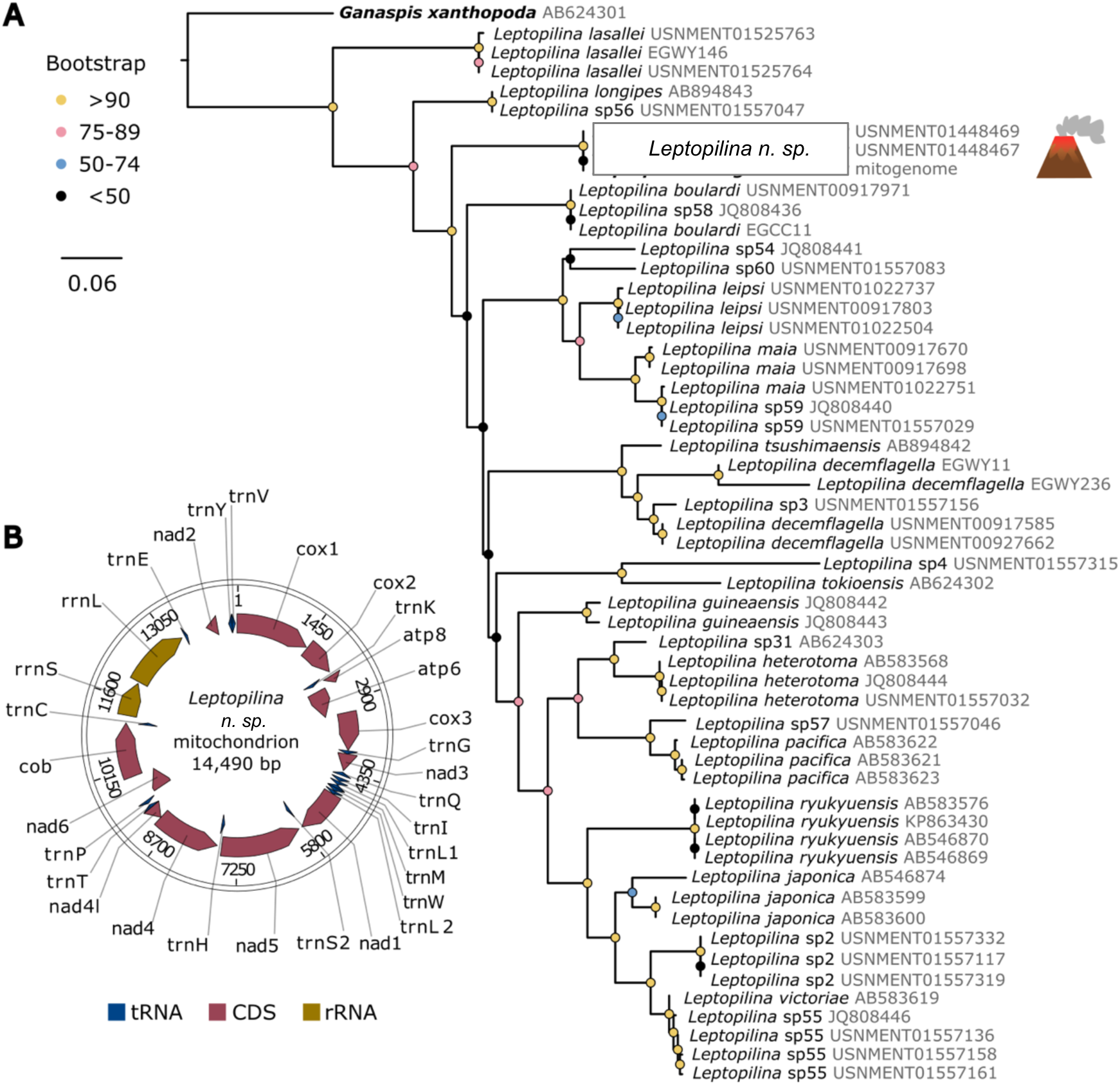
Mitochondrial phylogeny and genome of *Leptopilina n. sp.*. **(A)** Phylogenetic reconstruction of *Leptopilina,* based on 480 nucleotides of Cox1, aligned to maintain codon position relationships. Node colors indicate bootstrap support as indicated in legend. The three *Leptopilina n. sp.* sequences are identical. The two other *Leptopilina n. sp.* sequences are from previously barcoded STL6 specimens [3]. Sequences assigned to provisional species are indicated by ‘sp’ followed by a number, as per DROP database [3]. Accession numbers are in gray at the end of each leaf label. **(B)** Map of the assembled mitochondrial genome and annotations.

### Gene Annotation

Annotation of the repeat-masked *Leptopilina n. sp.* genome revealed 16,402 genes (Table 3), which is in-line with annotations of closely related species (Table 2). To better understand genome evolution, we clustered protein sequences (n=78,896 in total) of the four *Leptopilina* species plus outgroup species *Ganaspis kimorum* [6, 54, 56–58]. As expected with a small set of closely related species, we identified orthologs for most genes: 97.1% of the genes were clustered into 13,659 orthogroups, each of which contained 5.6 genes on average. 7,266 orthogroups contained genes from all five species, and of those, 6,389 orthogroups were strictly single-copy. 1,077 orthogroups were species-specific and only contained in-paralogs.

**Table 3.** Annotation metrics for the *Leptopilina n. sp.* genome.

| Metric | Value* |
| --- | --- |
| Number of genes | 16,402 |
| Number of mRNAs | 18,165 |
| Number of exons | 97,163 |
| Number of introns | 78,998 |
| Mean mRNAs per gene | 1.1 |
| Mean exons per mRNA | 5.3 |
| Total gene length | 158,352,048 bp |
| Longest gene | 235,629 bp |
| Mean gene length | 9,654 bp |
| Longest CDS | 76,353 bp |
| Mean CDS length | 1,541 bp |
| Longest exon | 16,278 bp |
| Mean exon length | 288 bp |

\*Base pairs = bp

### Repetitive elements

Given the larger than expected size of the *Leptopilina n. sp.* genome relative to other *Leptopilina* species, we analyzed the repeat content of the assembly using a *de novo* repeat identification pipeline. 66.81% of the nearly 1Gbp genome assembly was identified as repetitive (Figure 7, Supplementary Table S4). Retroelements and DNA transposons only constituted 9.57% and 5.09% of the length of the assembly, respectively. The majority of the repeat content (48.97% of the assembly) was categorized as unclassified repetitive elements. To determine if this large repetitive fraction was unique to *Leptopilina n. sp.* we ran the same *de novo* repeat identification pipeline on the three other *Leptopilina* species with genome assemblies available, plus an outgroup from a closely related genus, *Ganaspis kimorum* (Supplemental Table S4). Indeed, the other *Leptopilina* species were less repeat dense with 35.40%, 48.01%, and 57.27% repeat content for *L. clavipes, boulardi, and heterotoma,* respectively. Similar to *Leptopilina n. sp.*, the majority of these repetitive elements were unclassified. In fact, the overall size and makeup of the *Leptopilina n. sp.* genome was much more similar to the outgroup *Ganaspis kimorum* (Figure 7A,B). However, the *Ganaspis kimorum* and *Leptopilina n. sp.* genomes contained approximately double the amount of non-repetitive genome as compared to the other *Leptopilina* (black bars in Figure 7B). Thus, while *Leptopilina n. sp.* had the largest repetitive content in the genus (both by total length and percentage), repeat content alone does not completely explain the overall difference in genome size relative to the other congenerics.

**Figure 7.**
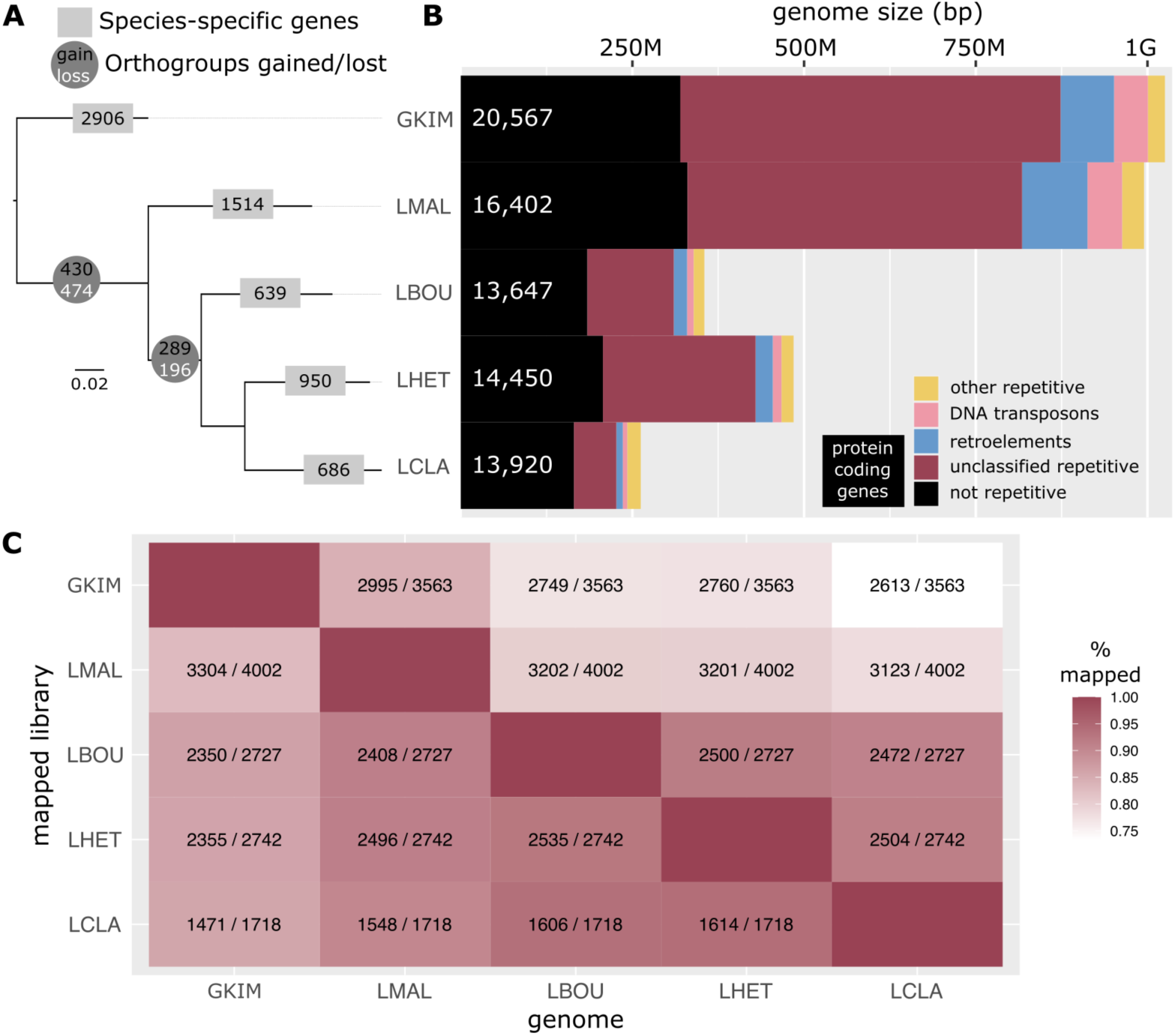
Evolution of *Leptopilina* genomes. **(A)** Species tree inferred by OrthoFinder, rooted on outgroup, *Ganaspis kimorum*. Phylogeny is annotated with (1) the number of species-specific genes (gray boxes), which include both singletons, and species-specific orthogroups, and (2) inferred gains and losses of orthogroups. “Gained” orthogroups are those that are unique to and present in all taxa within the clade, and “lost” orthogroups are present in all the outgroups, but none of the ingroups. **(B)** Genome content of each representative assembly. **(C)** All-verus-all comparison of repetitive elements. Libraries of the *de novo* identified repeats from each species were mapped to all other genomes, and the number of unique repeat families recovered in each comparison is indicated within each box in the heat map (number of families identified in “genome”/number of unique families in “library”). LMAL = *Leptopilina n. sp*.

To understand the large differences in repeat content across the *Leptopilina* genomes, we performed an all-versus-all analysis in which we mapped each species’ repeat library to all other species’ genomes (Figure 7C). Then we determined how many of the unique repeat families were present to infer if there was simply a difference in the number of repeat occurrences (e.g., family size), or, if entire repeat families had been gained or lost across the clade. The most repeat dense genomes, *Ganaspis kimorum* and *Leptopilina n. sp.*, contained 3,563 and 4,002 repeat families respectively, likely indicating a gain of repeats in the *Leptopilina* common ancestor. Accordingly, while 84.1% of the *Ganaspis kimorum* repeat families occur in the *Leptopilina n. sp.* genome, only 82.5% of the *Leptopilina n. sp.* repeat families occur in *Ganaspis kimorum*. There appears to have been a large loss of repeat families in the ancestor of the clade containing *L. boulardi*, *L. heterotoma*, and *L. clavipes*, and additional loss in the lineage leading to *L. clavipes*. These genomes contained 2,727, 2,742, and 1,718 families respectively, and between 88-91% of these families were found in *Leptopilina n. sp.*. Furthermore, while only 78-80% of the *Leptopilina n. sp.* repeats were identified in these other *Leptopilina* genomes, an even lower percentage of the *Ganaspis kimorum* repeats were present (73.3-77.5%). Finally, the comparisons within the clade of *L. boulardi*, *L. heterotoma*, and *L. clavipes*, indicate high levels of shared repeats (85.8-93.9%). All together, these data indicate that (1) the ancestor of the four *Leptopilina* species here acquired repetitive elements that are not present in *Ganaspis kimorum*, (2) many of the repeats have since been lost in *L. boulardi*, *L. heterotoma*, and especially in *L. clavipes*.

### *Wolbachia* Strain *w*Lmal

We identified a circular *Wolbachia* genome in our assembly, hereinafter strain “*w*Lmal” (Table 4). A single contig was sequenced at 344X coverage and blast results from blobtools revealed similarity to *Wolbachia* strain *w*Ha from *Drosophila simulans* across most of the contig. BUSCO genome completeness metrics were very high, with 99.2% of Rickettsiales BUSCOs present in single copy (Table 4). Phylogenetic reconstruction based on core single-copy orthologs revealed that *w*Lmal is closely related to *Drosophila* infecting *Wolbachia* strains (e.g., *w*Mel, *w*Ha*, w*Ri, *w*Au) in *Wolbachia* Supergroup A (Figure 8A). This is notable not only due to the host-parasitoid relationship of *Leptopilina* and *Drosophila*, but also because the parthenogenesis-inducing “*w*Lcla” strain from *Leptopilina clavipes* is quite divergent, and nested within a different major *Wolbachia* clade: Supergroup B [59, 60]. The size (∼1.2Mbp) and coding content of the *w*Lmal genome is quite typical of an insect-infecting *Wolbachia* strain [61, 62] (Figure 8B, Table 4). Two putative prophage regions were identified. The first (1,00,3471 bp – 1,051,238 bp) is partial and is likely remnants of a *Wolbachia* “WO” prophage. This prophage region contains some phage structural elements (primarily proteins for the tail) and what looks to be part of a “Eukaryotic Association Module” (EAM), a region associated with WO phages which often contains proteins for manipulating host reproduction and cell biology [63]. The second predicted prophage region (824,983 bp – 864,973 bp) appears much more degenerate and only has a few remaining open reading frames encoding for phage proteins.

**Figure 8.**
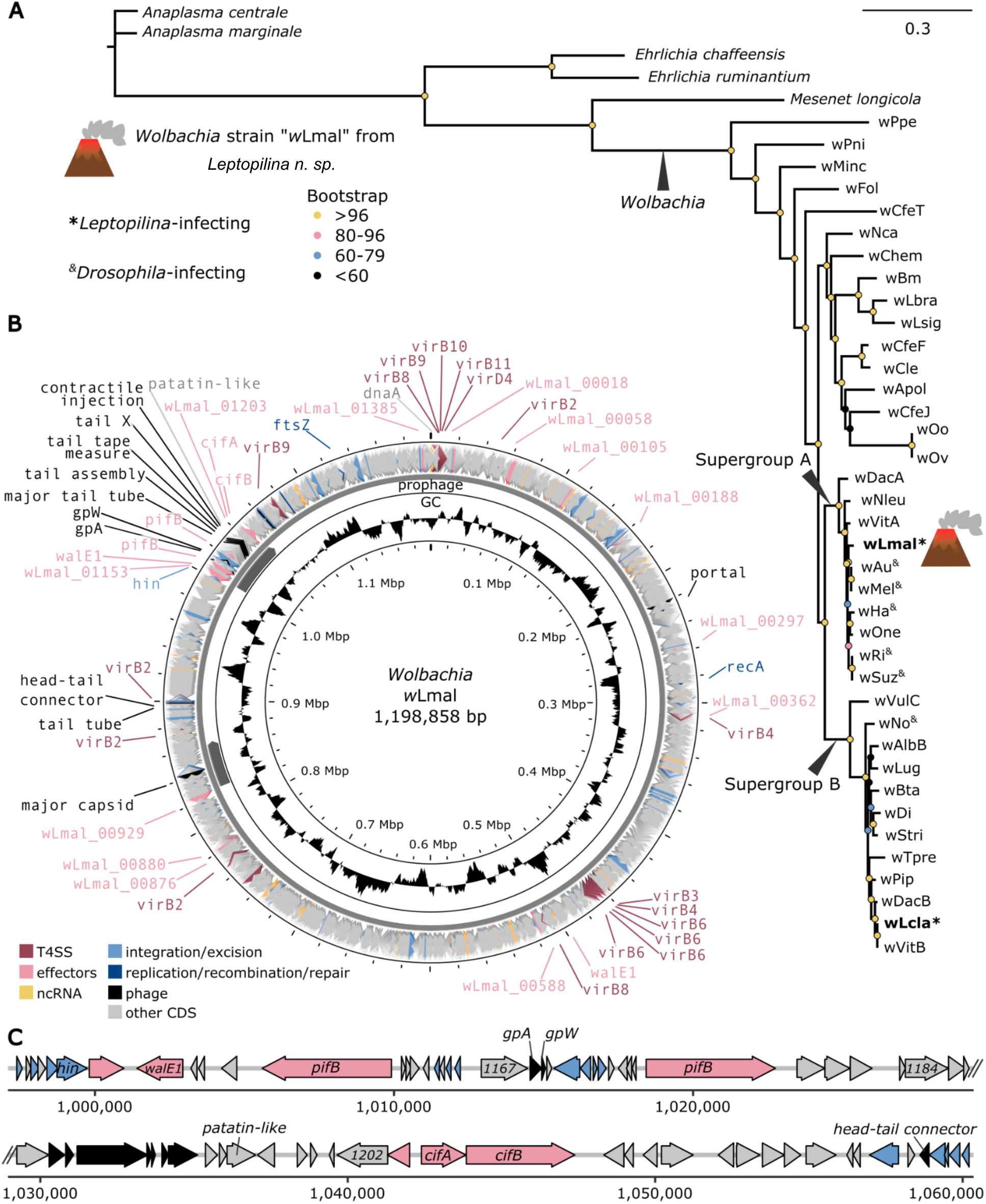
*Wolbachia* strain *w*Lmal. **(A)** Phylogenetic relationships of *Wolbachia* strains and Rickettsiales relatives based on 127 core single-copy orthologs. **(B)** Map of the complete *w*Lmal genome. Each concentric circle is annotated with a different set of genomic information. The innermost circle is annotated with a basepair marker, the second circle indicates GC content, the third circle indicates prophage regions identified by VirSorter, and the outermost circle shows the coding regions of the genome. Abbreviations: Type IV Secretion system (T4SS), noncoding RNA (ncRNA), coding sequences (CDS). Putative effector proteins are indicated with the gene name where possible, otherwise the wLmal CDS number is indicated and the *w*Mel homologs can be found in Supplemental Table S5. **(C)** Gene models in the prophage region containing remnant WO phage structural proteins and a suite of effector proteins homologs. Notably the two *pifB* coding sequences are present in an inverted repeat. Open reading frames labeled with numbers (e.g., 1167) are gene annotation IDs, provided to benchmark the region.

**Table 4.** *Wolbachia* strain *w*Lmal genome assembly and annotation.

| **Metric** | **wLmal** |
| --- | --- |
| **Contigs** | 1 |
| **Length (bp)** | 1,198,858 |
| **%GC** | 35.1 |
| **BUSCO\***<br>**(Rickettsiales)** | C:99.2%[S:99.2%,D:0.0%],<br>F:0.3%,M:0.5%,n:364 |
| **CDS** | 1,356 |
| **rRNAs** | 3 |
| **tRNAs** | 34 |

\*Standard BUSCO annotation: Complete BUSCOs (C) [Complete and single-copy BUSCOs (S), Complete and duplicated BUSCOs (D)], Fragmented BUSCOs (F), Missing BUSCOs (M), Total BUSCO groups searched (n)

We searched for the two recently identified putative parthenogenesis-inducing effector proteins, *pifA* and *pifB,* which are found in the parthenogenesis inducing strains *w*Lcla and *w*Tpre [59]. *pifA* is also present in the PI-*Wolbachia* infecting *Encarisa formosa* (designated *“piff”* in that strain) [64]. We identified two identical copies of *pifB*, but did not find evidence for *pifA* in *w*Lmal (Supplemental File S1). Closer inspection revealed that the *pifB* copies are in opposite orientations, and one copy of *pifB* is surrounded by insertion elements and is just downstream of a putative *hin* DNA invertase, likely from the WO phage. We also queried the *w*Lmal genome for a suite of other notable *Wolbachia* proteins and identified other putative or confirmed effector proteins. Surprisingly, these included the genes *cifA* and *cifB* (here, specifically “type I” or *cid*-type [65] which cause an entirely different host reproductive phenotype: the conditional sperm-egg incompatibility known as cytoplasmic incompatibility (CI).

Historical specimens of *Leptopilina n. sp*.

As noted above, our Saint Lucian specimens matched previously undescribed *Leptopilina* collected from the Caribbean in the 1980’s (specimens USNMENT01025775-USNMENT01025782). Specifically, those *Leptopilina* (n=16, all female) were collected in 1982 and 1983 from Grande Terre, Guadeloupe (∼270 km from our collection site on Saint Lucia), some of which were used to initiate a now extinct colony, “PB10-5”. The PB10-5 wasps were previously characterized as an undescribed species nested within an early-branching *Leptopilina* lineage, in the same clade as *Leptopilina longipes* [66, 67]. Our COI phylogenetic reconstruction of *Leptopilina* recapitulated these relationships (Figure 6A). In addition to all the collected Guadeloupe wasps being female, Yves Carton’s notes indicated “*[the PB10-5 colony] emerged only females: we can conclude that this species is parthenogenetic.*” In summary, our data indicate that *Leptopilina n. sp.* was previously collected ∼40 years ago on the neighboring island of Guadeloupe and also reproduced via thelytokous parthenogenesis.

## DISCUSSION

Our study represents a comprehensive description of *Leptopilina n. sp.*, the first recorded parasitoid of *dunni* group *Drosophila* from their native range. By integrating classic taxonomy and systematics approaches with genomics and experimental biology, we were able to generate a more comprehensive view of the biology and evolution of this new species. Furthermore, historical records and museum specimens enabled the discovery that asexually reproducing *Leptopilina n. sp.* have also been found on another island in the Antilles. This work represents one of a limited number of studies in which description of a new metazoan species is coupled with genome sequencing [68–72]. Indeed, this is the fifth such example in insects [73–76], wherein body size often precludes whole genome sequencing of holotypes.

Despite the interest in speciation patterns of the *dunni* group *Drosophila*, the proximal ecological or evolutionary mechanisms driving morphological and species diversity in this group remain unresolved. One study found elevated nonsynonymous substitution rates in the pigmentation gene *yellow* in two members of this group [77]. This is consistent with selection for pigmentation diversity, but no studies have yet identified the selective agent responsible for this pattern. Importantly, melanization is a process in many insect lineages that is critical in both the development of cuticle pigmentation as well as in immune responses that occur during encapsulation of foreign pathogens [78]. In *Drosophila*, melanization is a well-studied response to infection by eukaryotic parasites (including *Leptopilina*) and is linked to the same genetic pathways as pigmentation production [79–81]. In one study in *Drosophila falleni*, selection for a reduced number of abdominal spots resulted in higher infection rates by a parasitic nematode [82]. In other insects there are similar links between the levels or patterns of cuticular melanization and resistance to parasites [83–88]. As such, it has been postulated that *Drosophila* abdominal coloration may vary in some systems primarily due to selection by parasitism [89, 90], but direct evidence for this is lacking.

Since adult flies of the *D. antillea* species are among the more highly pigmented flies of the *dunni* group, we were interested in the possibility that they might be particularly resistant to infection by endemic parasitoids. However, we instead found the opposite. *Leptopilina n. sp.* wasps are highly virulent on *D. antillea* hosts. This observation could indicate that *Leptopilina n. sp.* is a relatively recent colonizer of Saint Lucia and that *D. antillea* has not had sufficient time to evolve resistance. In contrast, *Leptopilina n. sp.* was less effective at using *D. willistoni* as a host. Our findings open the door to future studies to explore the potential mechanism of *Leptopilina n. sp.* virulence on flies of the *dunni* group as well as the potential mechanisms of resistance for *D. willistoni.* It will also be of interest to test the virulence of *Leptopilina n. sp.* on other species from the *dunni* group as well as how virulence tracks with island of origin of the wasp. For example, are *Leptopilina n. sp.* wasps as effective when infecting *dunni* group flies that are native to other islands of the Antilles chain? Are *Leptopilina n. sp.* wasps that originate from Guadeloupe as virulent on *D. antillea*? Indeed, the island of Guadeloupe where Carton collected *Leptopilina n. sp.* is home to the dunni group fly *Drosophila arawakana.* It would be of interest to know how the virulence of Guadeloupian wasps relates to Saint Lucian wasps on fly species from the various islands.

The island biogeography of *Leptopilina n. sp.* may also play a role in the evolution of their reproductive mode. Our antibiotic curing experiments indicated that our *Leptopilina n. sp.* colony derived from Saint Lucia is unable to revert to sexual reproduction. The inability to restore sex is a strong indicator of a so-called “virginity mutation”, which is well described in several other wasp species with symbiont-mediated parthenogenesis [57, 91–96]. There are multiple hypotheses for the evolutionary pressures that lead to virginity mutations [48, 91, 97, 98], but in brief, they are mutations that result in the decay or loss of sexual function, to the point that the animal becomes reliant upon the PI symbiont for the production of female offspring. Related to this, it is notable that the colony of *Leptopilina n. sp.* derived from Guadeloupe collections in the 1980s were all female. However, we do not know if this PB10-5 colony was (a) infected with *Wolbachia*, (b) asexual because of *Wolbachia*, or (c) obligately asexual (i.e., unable to revert to a sexual mode). Indeed, some other populations of parasitic wasps with PI *Wolbachia* have significant within-species polymorphism and vary in which *Wolbachia* strain they have, if they are asexually reproducing, and if they have a so-called virginity mutation [48, 99]. Thus, it may be that populations of *Leptopilina n. sp.* on different islands vary in whether they are sexually reproducing, and if asexual reproduction is fixed or reversible. Notably, asexual reproduction across many taxa appears to be more common on islands [100, 101], and thus *Leptopilina n. sp.* in the Greater Antilles may provide a tractable system to understand the evolution of reproductive mode in the context of biogeography.

While there are many other examples of species with virginity mutations, our findings are quite unusual in that the *Wolbachia* genome here also contains what appears to be functional genes for inducing cytoplasmic incompatibility (CI). Paradoxically, the CI phenotype only manifests with sexual reproduction: males with CI *Wolbachia*-modified sperm cause embryonic lethality in embryos derived from females without a compatible *Wolbachia* strain [65]. We have several hypotheses for why these *Wolbachia* encode for CI loci despite inducing parthenogenesis. First, the *cif* loci may have undergone some form of neofunctionalization: perhaps these proteins are no longer involved in sperm-egg incompatibility as in CI but are impacting other host processes. Alternatively, it might be that the transition to parthenogenesis is more recent, and CI is no longer under selection. Perhaps the *cifs* are en route to be eliminated from the genome in the future. “Recent” is of course difficult to determine quantitatively, especially in island systems where there are often strong bottlenecks that can drive rapid evolution [102]. Finally, it might be that if the PI phenotype has lower penetrance and occasional males are produced. These males might mate with incompatible conspecifics (that either have no *Wolbachia* or an incompatible strain) and induce CI, perhaps reducing competition for the asexual matriline. However, we see two issues with this scenario. First, lower penetrance of PI phenotypes is typically due to the loss of *Wolbachia* (e.g., due to higher temperatures, or in the lab, antibiotics), which leads to the production of males. In this case, the occasional males produced by PI mothers might not have sufficient *Wolbachia* to induce CI. Second, because the PI females transmitting *Wolbachia* to the next generation would never need to rescue CI, there would be no selection pressure to maintain CI function.

Despite the peculiarity of a PI *Wolbachia* strain encoding for CI genes, all four loci associated with host reproductive manipulation (*cifA, cifB,* and two copies of *pifB*) are found in the EAM of WO: a classic signature of *Wolbachia* genomes and reproductive effector proteins. The multiple identical *pifB* copies, genes for CI in combination with a clear *Wolbachia*-mediated PI phenotype, and evidence for multiple prophage infections all point towards recent dynamism in the *w*Lmal genome. Many *Wolbachia* genes mediating host reproductive switches are horizontally acquired from other *Wolbachia* strains, likely due to the WO phage [63, 103]. For this reason, it was not surprising that the two PI *Wolbachia* strains from *Leptopilina* both encoded for PifB proteins, despite the strains belonging to different clades. In contrast, we did not find any evidence in *w*Lmal for the PifA protein, a *transformer* mimic that interacts with the sexual differentiation pathway [59, 64]. Importantly however, we do not know the exact mechanism of parthenogenesis induction employed by *w*Lmal (e.g., one versus two step, apomixis versus gamete duplication, etc [104]). And, given the diversity of PI mechanisms found across *Wolbachia* [48] it is quite possible that *w*Lcla and *w*Lmal are not using precisely the same toolset. Closer inspection of meiosis and early embryonic mitoses in *Leptopilina n. sp.* will be key to understanding how *w*Lmal causes PI, and what the role of PifB, or other proteins, might be in this process. Altogether, the description of *Leptopilina n. sp.* and its reproductive biology will enable future explorations of host-parasitoid and host-symbiont interactions, especially in the context of speciation.

In summary, this study provides a clear example of how descriptive work in under-studied species can lead to novel discoveries across a broad set of biological disciplines. Insects generally, but especially parasitic Hymenoptera, are under described [5]. Furthermore, of the described parasitic hymenopterans, a significant majority lack biological information such as host associations and ecology. Here, identification of a new species of parasitoid has provided a unique view into the intricate co-evolution of host insect, parasitoid, and *Wolbachia* symbiont, with dynamic interactions between genomic evolution, virulence, and reproductive ecology. We hope our work inspires more integration of descriptions of new species coupled with investigations into novel aspects of their biology.

## DATA AVAILABILITY

BioProject: PRJNA1150626

Supplemental File S1 contains:

Supplemental Text

Figure S1. Western Blot for *Wolbachia* detection.

Figure S2. Draft assembly snail plot and blob plot.

Supplemental Tables are provided in an excel file where each sheet corresponds to a different table number:

Table S1. Passed reads statistics.

Table S2. All draft assembly contigs statistics, including BlobTools annotations.

Table S3. Purged contigs.

Table S4. Repeat annotations.

Table S5. *Wolbachia* protein queries.

Table S6. wLmal phage annotation.

## ACKNOWLEDGEMENTS

The research expedition of the sailing vessel Sea Salt was conceived in conversations with W. Daniel Tracey Sr. (now deceased), and we dedicate this study to his memory. We are very thankful to Amalia Rowan the first mate on the Sea Salt without whose assistance our collection expedition could not have taken place. We are thankful for Dr. Hannah Romain (Saint Lucia Ministry of Agriculture) for granting permission to collect insects on Saint Lucia. We are also thankful to Michael Gerth for making the WHOP pipeline available and to Keith Hopper for sharing the *Ganaspis* assembly prior to its release. Research reported in this publication was supported by the National Institute of General Medical Sciences of the National Institutes of Health under award number R35GM150991 to ARIL, the National Institute of Food and Agriculture Predoctoral Fellowship to LCF (NIFA 2023-67034-40496), and the Linda and Jack Gill Institute for Neuroscience Research Stipend to WDT. Mention of trade names or commercial products in this publication is solely for the purpose of providing specific information and does not imply recommendation or endorsement by the USDA. USDA is an equal opportunity provider and employer.

## CITATIONS

1. Karvonen A, Seehausen O. The role of parasitism in adaptive radiations—when might parasites promote and when might they constrain ecological speciation? International Journal of Ecology. 2012;2012(1):280169.

2. Nyman T, Bokma F, Kopelke J-P. Reciprocal diversification in a complex plant-herbivore-parasitoid food web. BMC Biol. 2007;5:1–9.

3. Lue CH, Buffington ML, Scheffer S, Lewis M, Elliott TA, Lindsey AR, et al. DROP: Molecular voucher database for identification of *Drosophila* parasitoids. Molecular Ecology Resources. 2021;21(7):2437–54.

4. Saunders TE, Ward DF. Variation in the diversity and richness of parasitoid wasps based on sampling effort. PeerJ. 2018;6:e4642.

5. Forbes AA, Bagley RK, Beer MA, Hippee AC, Widmayer HA. Quantifying the unquantifiable: why Hymenoptera, not Coleoptera, is the most speciose animal order. BMC Ecol. 2018;18:1–11.

6. Sosa-Calvo J, Forshage M, Buffington ML. Circumscription of the Ganaspis brasiliensis (Ihering, 1905) species complex (Hymenoptera, Figitidae), and the description of two new species parasitizing the spotted wing drosophila, Drosophila suzukii Matsumura, 1931 (Diptera, Drosophilidae). J Hymenoptera Res. 2024;97:441-70.

7. Coyne JA, Orr HA. Speciation: a catalogue and critique of species concepts. Philosophy of Biology: An Anthology. 2004:272–92.

8. Carson HL, Templeton AR. Genetic revolutions in relation to speciation phenomena: the founding of new populations. Annu Rev Ecol Syst. 1984:97–131.

9. Kaneshiro K, Gillespie R, Carson H. Chromosomes and male genitalia of Hawaiian *Drosophila*: tools for interpreting phylogeny and geography. Hawaiian Biogeography: Evolution on a hot spot archipelago. 1995:57–71.

10. O’Grady P, DeSalle R. Hawaiian *Drosophila* as an evolutionary model clade: days of future past. Bioessays. 2018;40(5):1700246.

11. Dworkin I, Jones CD. Genetic changes accompanying the evolution of host specialization in *Drosophila sechellia*. Genetics. 2009;181(2):721–36.

12. Hollocher H. Island hopping in *Drosophila*: patterns and processes. Philosophical Transactions of the Royal Society of London Series B: Biological Sciences. 1996;351(1341):735-43.

13. Heed WB, Krishnamurthy NB. Genetic studies on the cardini group of *Drosophila* in the West Indies. Univ Texas Publ. 1959;5914:155–79.

14. Hollocher H, Hatcher JL, Dyreson EG. Genetic and developmental analysis of abdominal pigmentation differences across species in the *Drosophila dunni* subgroup. Evolution. 2000;54(6):2057–71.

15. Lue C-H, Driskell AC, Leips J, Buffington ML. Review of the genus *Leptopilina* (Hymenoptera, Cynipoidea, Figitidae, Eucoilinae) from the Eastern United States, including three newly described species. J Hymenoptera Res. 2016;(53).

16. Stacconi MVR, Buffington M, Daane KM, Dalton DT, Grassi A, Kaçar G, et al. Host stage preference, efficacy and fecundity of parasitoids attacking *Drosophila suzukii* in newly invaded areas. Biol Control. 2015;84:28–35.

17. Abram PK, Franklin MT, Hueppelsheuser T, Carrillo J, Grove E, Eraso P, et al. Adventive larval parasitoids reconstruct their close association with spotted-wing drosophila in the invaded North American range. Environ Entomol. 2022;51(4):670–8.

18. Newton IL, Savytskyy O, Sheehan KB. Wolbachia utilize host actin for efficient maternal transmission in Drosophila melanogaster. PLoS Path. 2015;11(4):e1004798. doi: 10.1371/journal.ppat.1004798. PubMed PMID: 25906062; PubMed Central PMCID: PMCPMC4408098.

19. Lindsey AR, Tennessen JM, Gelaw MA, Jones MW, Parish AJ, Newton IL, et al. The intracellular symbiont Wolbachia alters Drosophila development and metabolism to buffer against nutritional stress. bioRxiv. 2023.

20. Fricke LC, Lindsey AR. Examining *Wolbachia*-Induced Parthenogenesis in Hymenoptera. Wolbachia: Methods and Protocols: Springer; 2024. p. 55–68.

21. Shen W, Le S, Li Y, Hu F. SeqKit: a cross-platform and ultrafast toolkit for FASTA/Q file manipulation. PLoS One. 2016;11(10):e0163962.

22. Kolmogorov M, Yuan J, Lin Y, Pevzner PA. Assembly of long, error-prone reads using repeat graphs. Nat Biotechnol. 2019;37(5):540–6.

23. Roach MJ, Schmidt SA, Borneman AR. Purge Haplotigs: allelic contig reassignment for third-gen diploid genome assemblies. BMC Bioinformatics. 2018;19(1):1–10.

24. Li H. Aligning sequence reads, clone sequences and assembly contigs with BWA-MEM. arXiv preprint arXiv:13033997. 2013.

25. Li H, Handsaker B, Wysoker A, Fennell T, Ruan J, Homer N, et al. The Sequence Alignment/Map format and SAMtools. Bioinformatics. 2009;25(16):2078–9. doi: 10.1093/bioinformatics/btp352.

26. Challis R, Richards E, Rajan J, Cochrane G, Blaxter M. BlobToolKit–interactive quality assessment of genome assemblies. G3: Genes, Genomes, Genetics. 2020;10(4):1361–74. PubMed Central PMCID: PMCPMC7144090.

27. Simão FA, Waterhouse RM, Ioannidis P, Kriventseva EV, Zdobnov EM. BUSCO: Assessing genome assembly and annotation completeness with single-copy orthologs. Bioinformatics. 2015. doi: 10.1093/bioinformatics/btv351.

28. Bernt M, Donath A, Jühling F, Externbrink F, Florentz C, Fritzsch G, et al. MITOS: improved de novo metazoan mitochondrial genome annotation. Mol Phylogen Evol. 2013;69(2):313–9.

29. Wanner N, Larsen PA, McLain A, Faulk C. The mitochondrial genome and Epigenome of the Golden lion Tamarin from fecal DNA using Nanopore adaptive sequencing. BMC Genomics. 2021;22:1–11.

30. Katoh K, Standley DM. MAFFT multiple sequence alignment software version 7: improvements in performance and usability. Mol Biol Evol. 2013;30(4):772–80. doi: 10.1093/molbev/mst010.

31. Nguyen L-T, Schmidt HA, Von Haeseler A, Minh BQ. IQ-TREE: a fast and effective stochastic algorithm for estimating maximum-likelihood phylogenies. Mol Biol Evol. 2015;32(1):268–74.

32. Flynn JM, Hubley R, Goubert C, Rosen J, Clark AG, Feschotte C, et al. RepeatModeler2 for automated genomic discovery of transposable element families. Proc Natl Acad Sci. 2020;117(17):9451–7.

33. Storer J, Hubley R, Rosen J, Wheeler TJ, Smit AF. The Dfam community resource of transposable element families, sequence models, and genome annotations. Mobile DNA. 2021;12:1–14.

34. Keilwagen J, Hartung F, Grau J. GeMoMa: homology-based gene prediction utilizing intron position conservation and RNA-seq data. Gene Prediction: Methods and Protocols. 2019:161–77.

35. Dobin A, Gingeras TR. Mapping RNA-seq reads with STAR. Current Protocols in Bioinformatics. 2015;51(1):11.4. 1-.4. 9.

36. Emms DM, Kelly S. OrthoFinder: phylogenetic orthology inference for comparative genomics. Genome Biol. 2019;20:1–14.

37. Gerth M. WHOP (v0.1). Zenodo. 2023. doi: 10.5281/zenodo.8343898.

38. Seemann T. Prokka: rapid prokaryotic genome annotation. Bioinformatics. 2014;30(14):2068–9.

39. Bruen T, Bruen T. PhiPack: PHI test and other tests of recombination. McGill University, Montreal, Quebec. 2005:1-8.

40. Grant JR, Enns E, Marinier E, Mandal A, Herman EK, Chen C-y, et al. Proksee: in-depth characterization and visualization of bacterial genomes. Nucleic Acids Res. 2023;51(W1):W484–W92.

41. Guo J, Bolduc B, Zayed AA, Varsani A, Dominguez-Huerta G, Delmont TO, et al. VirSorter2: a multi-classifier, expert-guided approach to detect diverse DNA and RNA viruses. Microbiome. 2021;9:1–13.

42. Starikova EV, Tikhonova PO, Prianichnikov NA, Rands CM, Zdobnov EM, Ilina EN, et al. Phigaro: high-throughput prophage sequence annotation. Bioinformatics. 2020;36(12):3882–4.

43. Brown CL, Mullet J, Hindi F, Stoll JE, Gupta S, Choi M, et al. mobileOG-db: a manually curated database of protein families mediating the life cycle of bacterial mobile genetic elements. Appl Environ Microbiol. 2022;88(18):e00991–22.

44. R Core Team. R: A language and environment for statistical computing. R Foundation for Statistical Computing, Vienna, Austria: URL http://www.R-project.org/; 2014.

45. Frerebeau N, Lebrun B, Arel-Bundock V. khroma: Colour Schemes for Scientific Data Visualization. 2019.

46. Wachi N, Nomano FY, Mitsui H, Kasuya N, Kimura MT. Taxonomy and evolution of putative thelytokous species of *Leptopilina* (Hymenoptera: Figitidae) from Japan, with description of two new species. Entomol Sci. 2015;18(1):41–54.

47. Chen F, Schenkel M, Geuverink E, van de Zande L, Beukeboom LW. Absence of complementary sex determination in two Leptopilina species (Figitidae, Hymenoptera) and a reconsideration of its incompatibility with endosymbiont-induced thelytoky. Insect Sci. 2022;29(3):900–14.

48. Ma WJ, Schwander T. Patterns and mechanisms in instances of endosymbiont-induced parthenogenesis. J Evol Biol. 2017;30(5):868–88.

49. Jervis MA, Heimpel GE, Ferns PN, Harvey JA, Kidd NA. Life-history strategies in parasitoid wasps: a comparative analysis of ‘ovigeny’. J Anim Ecol. 2001;70(3):442–58.

50. Doremus MR, Stouthamer CM, Kelly SE, Schmitz-Esser S, Hunter MS. Quality over quantity: unraveling the contributions to cytoplasmic incompatibility caused by two coinfecting *Cardinium* symbionts. Heredity. 2022;128(3):187–95.

51. Duron O, Bouchon D, Boutin S, Bellamy L, Zhou L, Engelstädter J, et al. The diversity of reproductive parasites among arthropods: *Wolbachia* do not walk alone. BMC Biol. 2008;6(1):1–12.

52. Kageyama D, Narita S, Watanabe M. Insect sex determination manipulated by their endosymbionts: incidences, mechanisms and implications. Insects. 2012;3(1):161–99.

53. Perlmutter JI, Bordenstein SR. Microorganisms in the reproductive tissues of arthropods. Nat Rev Micro. 2020;18(2):97–111.

54. Hopper KR, Wang X, Kenis M, Seehausen ML, Abram PK, Daane KM, et al. Genome divergence and reproductive incompatibility among populations of Ganaspis near brasiliensis. G3: Genes, Genomes, Genetics. 2024:jkae090.

55. Cameron SL. Insect mitochondrial genomics: implications for evolution and phylogeny. Annu Rev Entomol. 2014;59(1):95–117.

56. Khan S, Sowpati DT, Srinivasan A, Soujanya M, Mishra RK. Long-read genome sequencing and assembly of Leptopilina boulardi: a specialist Drosophila parasitoid. G3: Genes, Genomes, Genetics. 2020;10(5):1485–94.

57. Kraaijeveld K, Anvar SY, Frank J, Schmitz A, Bast J, Wilbrandt J, et al. Decay of sexual trait genes in an asexual parasitoid wasp. Genome Biol Evol. 2016;8(12):3685–95.

58. Wey B, Heavner ME, Wittmeyer KT, Briese T, Hopper KR, Govind S. Immune suppressive extracellular vesicle proteins of Leptopilina heterotoma are encoded in the wasp genome. G3: Genes, Genomes, Genetics. 2020;10(1):1–12.

59. Fricke LC, Lindsey AR. Identification of parthenogenesis-inducing effector proteins in *Wolbachia*. Genome Biol Evol. 2024:evae036.

60. Lindsey ARI, Werren JH, Richards S, Stouthamer R. Comparative genomics of a parthenogenesis-inducing Wolbachia symbiont. G3: Genes|Genomes|Genetics. 2016;6(7):2113–23. Epub 2016/05/20. doi: 10.1534/g3.116.028449. PubMed PMID: 27194801; PubMed Central PMCID: PMCPMC4938664.

61. Kaur R, Shropshire JD, Cross KL, Leigh B, Mansueto AJ, Stewart V, et al. Living in the endosymbiotic world of *Wolbachia*: A centennial review. Cell Host & Microbe. 2021;29(6):879–93. PubMed Central PMCID: PMC8192442.

62. Lindsey ARI. Sensing, Signaling, and Secretion: A review and analysis of systems for regulating host interaction in Wolbachia. Genes. 2020;11(7):813. PubMed Central PMCID: PMC7397232.

63. Bordenstein SR, Bordenstein SR. Eukaryotic association module in phage WO genomes from *Wolbachia*. Nat Commun. 2016;7.

64. Li C, Li C-Q, Chen Z-B, Liu B-Q, Sun X, Wei K-H, et al. *Wolbachia* symbionts control sex in a parasitoid wasp using a horizontally acquired gene. Curr Biol. 2024.

65. Beckmann JF, Bonneau M, Chen H, Hochstrasser M, Poinsot D, Merçot H, et al. The toxin– antidote model of cytoplasmic incompatibility: genetics and evolutionary implications. Trends Genet. 2019;35(3):175–85. PubMed Central PMCID: PMC6519454

66. Schilthuizen M, Nordlander G, Stouthamer R, van Alphen J. Morphological and molecular phylogenetics in the genus *Leptopilina* (Hymenoptera: Cynipoidea: Eucoilidae). Syst Entomol. 1998;23(3):253–64.

67. Vet LE, van Alphen JJ. A comparative functional approach to the host detection behaviour of parasitic wasps. 1. A qualitative study on Eucoilidae and Alysiinae. Oikos. 1985:478–86.

68. Kanzaki N, Tsai IJ, Tanaka R, Hunt VL, Liu D, Tsuyama K, et al. Biology and genome of a newly discovered sibling species of *Caenorhabditis elegans*. Nat Commun. 2018;9(1):3216.

69. Koehler G, Khaing KPP, Than NL, Baranski D, Schell T, Greve C, et al. A new genus and species of mud snake from Myanmar (Reptilia, Squamata, Homalopsidae). Zootaxa. 2021;4915(3):zootaxa. 4915.3. 1-zootaxa. 3. 1.

70. Köhler G, Vargas J, Than NL, Schell T, Janke A, Pauls SU, et al. A taxonomic revision of the genus Phrynoglossus in Indochina with the description of a new species and comments on the classification within Occidozyginae (Amphibia, Anura, Dicroglossidae). Vertebrate Zoology. 2021;71:1-26.

71. Köhler G, Zwitzers B, Than NL, Gupta DK, Janke A, Pauls SU, et al. Bioacoustics reveal hidden diversity in frogs: Two new species of the genus Limnonectes from Myanmar (Amphibia, Anura, Dicroglossidae). Diversity. 2021;13(9):399.

72. Sullivan JP, Hopkins CD, Pirro S, Peterson R, Chakona A, Mutizwa TI, et al. Mitogenome recovered from a 19th Century holotype by shotgun sequencing supplies a generic name for an orphaned clade of African weakly electric fishes (Osteoglossomorpha Mormyridae). ZooKeys. 2022;1129:163.

73. Brandão-Dias PF, Zhang YM, Pirro S, Vinson CC, Weinersmith KL, Ward AK, et al. Describing biodiversity in the genomics era: A new species of Nearctic Cynipidae gall wasp and its genome. Syst Entomol. 2022;47(1):94–112.

74. Heckenhauer J, Razuri-Gonzales E, Mwangi FN, Schneider J, Pauls SU. Holotype sequencing of Silvatares holzenthali Rázuri-Gonzales, Ngera & Pauls, 2022 (Trichoptera, Pisuliidae). ZooKeys. 2023;1159:1.

75. Pohl H, Niehuis O, Gloyna K, Misof B, Beutel RG. A new species of *Mengenilla* (Insecta, Strepsiptera) from Tunisia. ZooKeys. 2012;(198):79.

76. Rázuri-Gonzales E, Graf W, Heckenhauer J, Schneider JV, Pauls SU. A new species of Rhyacophila Pictet, 1834 (Trichoptera, Rhyacophilidae) from Corsica with the genomic characterization of the holotype. ZooKeys. 2024;1218:295.

77. Wilder J, Dyreson E, O’Neill R, Spangler M, Gupta R, Wilder A, et al. Contrasting modes of natural selection acting on pigmentation genes in the *Drosophila dunni* subgroup. Journal of Experimental Zoology Part B: Molecular and Developmental Evolution. 2004;302(5):469–82.

78. Sugumaran M, Barek H. Critical analysis of the melanogenic pathway in insects and higher animals. International Journal of Molecular Sciences. 2016;17(10):1753.

79. Dolezal T. How to eliminate pathogen without killing oneself? Immunometabolism of encapsulation and melanization in *Drosophila*. Frontiers in Immunology. 2023;14:1330312.

80. Carton Y, Poirié M, Nappi AJ. Insect immune resistance to parasitoids. Insect Sci. 2008;15(1):67–87.

81. Russo J, Dupas S, Frey F, Carton Y, Brehelin M. Insect immunity: early events in the encapsulation process of parasitoid (*Leptopilina boulardi*) eggs in resistant and susceptible strains of *Drosophila*. Parasitology. 1996;112(1):135–42.

82. Dombeck I, Jaenike J. Ecological genetics of abdominal pigmentation in *Drosophila falleni*: a pleiotropic link to nematode parasitism. Evolution. 2004;58(3):587–96.

83. Siva–Jothy MT. A mechanistic link between parasite resistance and expression of a sexually selected trait in a damselfly. Proceedings of the Royal Society of London Series B: Biological Sciences. 2000;267(1461):2523-7.

84. Wilson K, Cotter SC, Reeson AF, Pell JK. Melanism and disease resistance in insects. Ecol Lett. 2001;4(6):637–49.

85. Tan S, Wang Y, Liu P, Ge Y, Li A, Xing Y, et al. Increase of albinistic hosts caused by gut parasites promotes self-transmission. Frontiers in Microbiology. 2018;9:1525.

86. Lindsey E, Altizer S. Sex differences in immune defenses and response to parasitism in monarch butterflies. Evol Ecol. 2009;23:607–20.

87. Krams I, Burghardt GM, Krams R, Trakimas G, Kaasik A, Luoto S, et al. A dark cuticle allows higher investment in immunity, longevity and fecundity in a beetle upon a simulated parasite attack. Oecologia. 2016;182:99–109.

88. Dubovskiy I, Whitten M, Kryukov V, Yaroslavtseva O, Grizanova E, Greig C, et al. More than a colour change: insect melanism, disease resistance and fecundity. Proc R Soc Lond B. 2013;280(1763):20130584.

89. Wittkopp PJ, Beldade P, editors. Development and evolution of insect pigmentation: genetic mechanisms and the potential consequences of pleiotropy. Semin Cell Dev Biol; 2009: Elsevier.

90. Kohler LJ, Carton Y, Mastore M, Nappi AJ. Parasite suppression of the oxidations of eumelanin precursors in *Drosophila melanogaster*. Archives of Insect Biochemistry and Physiology: Published in Collaboration with the Entomological Society of America. 2007;66(2):64–75.

91. Stouthamer R, Russell JE, Vavre F, Nunney L. Intragenomic conflict in populations infected by Parthenogenesis Inducing *Wolbachia* ends with irreversible loss of sexual reproduction. BMC Evol Biol. 2010;10:12. doi: 10.1186/1471-2148-10-229. PubMed PMID: WOS:000282746700002.

92. Ma W-J, Pannebakker BA, Li X, Geuverink E, Anvar SY, Veltsos P, et al. A single QTL with large effect is associated with female functional virginity in an asexual parasitoid wasp. Mol Ecol. 2021;30(9):1979–92. doi: 10.1111/mec.15863.

93. Ma WJ, Pannebakker BA, Beukeboom LW, Schwander T, van de Zande L. Genetics of decayed sexual traits in a parasitoid wasp with endosymbiont-induced asexuality. Heredity. 2014;113(5):424–31. doi: 10.1038/hdy.2014.43. PubMed PMID: 24781809; PubMed Central PMCID: PMC4220718.

94. Russell JE, Stouthamer R. The genetics and evolution of obligate reproductive parasitism in *Trichogramma pretiosum* infected with parthenogenesis-inducing *Wolbachia*. Heredity. 2011;106(1):58–67. doi: 10.1038/hdy.2010.48. PubMed PMID: WOS:000285336100007.

95. Gottlieb Y, Zchori-Fein E. Irreversible thelytokous reproduction in *Muscidifurax uniraptor*. Entomol Exp Appl. 2001;100(3):271–8.

96. Stouthamer R, Mak F. Influence of antibiotics on the offspring production of the *Wolbachia*-infected parthenogenetic parasitoid *Encarsia formosa*. J Invertebr Pathol. 2002;80(1):41–5.

97. van der Kooi CJ, Schwander T. On the fate of sexual traits under asexuality. Biol Rev Camb Philos Soc. 2014;89(4):805–19. doi: 10.1111/brv.12078. PubMed PMID: 24443922.

98. Neiman M, Sharbel TF, Schwander T. Genetic causes of transitions from sexual reproduction to asexuality in plants and animals. J Evol Biol. 2014;27(7):1346–59. doi: 10.1111/jeb.12357. PubMed PMID: 24666600.

99. van der Kooi CJ, Matthey-Doret C, Schwander T. Evolution and comparative ecology of parthenogenesis in haplodiploid arthropods. Evolution Letters. 2017;1(6):304–16.

100. Vrijenhoek RC, Parker ED. Geographical parthenogenesis: general purpose genotypes and frozen niche variation. Lost sex: the evolutionary biology of parthenogenesis. 2009:99–131.

101. Hörandl E. Geographical parthenogenesis: opportunities for asexuality. Lost sex: The evolutionary biology of parthenogenesis. 2009:161–86.

102. Gillespie RG. Island time and the interplay between ecology and evolution in species diversification. Evolutionary Applications. 2016;9(1):53–73.

103. Lindsey ARI, Rice DW, Bordenstein SR, Brooks AW, Bordenstein SR, Newton ILG. Evolutionary genetics of cytoplasmic incompatibility genes *cifA* and *cifB* in prophage WO of *Wolbachia*. Genome Biol Evol. 2018;10(2):434–51. doi: 10.1093/gbe/evy012. PubMed Central PMCID: PMCPMC5793819.

104. Verhulst EC, Pannebakker BA, Geuverink E. Variation in sex determination mechanisms may constrain parthenogenesis-induction by endosymbionts in haplodiploid systems. Current Opinion in Insect Science. 2023:101023.

