## Supplemental Text and Figures for "Genomics and reproductive biology of *Leptopilina n. sp.* Buffington, Lue, Davis & Tracey sp. nov. (Hymenoptera: Figitidae): An asexual parasitoid of Caribbean *Drosophila*"

### RESULTS

#### Curating microbial contamination in the draft assembly

There were three short, low-coverage contigs that we manually inspected due to "best species" annotations as *Rickettsia* via blobtools (Supplemental Table S3). Further inspection revealed that these three contigs shared similarity with other insect sequences (including *Leptopilina*), and only short regions of these contigs contained matches to *Rickettsia* (≤ 10% of the contig length). Therefore, we are confident these contigs do not indicate an active *Rickettsia* infection in the wasps, and it is likely the contigs are not truly rickettsial in origin.

#### A spurious *pifB* containing contig is an assembly artifact

When searching for *pif* loci in the draft assembly, we found a single short additional contig that contained another identical copy of *pifB*. However, this appears to be an assembly artifact: the 6,631 bp accessory contig (contig_2041) matches the regions in the *w*Lmal genome surrounding the two *pifB* ORFs. In both cases, the sequence from 1,831 bp upstream of each *pifB* ORF to 477 bp downstream of each *pifB* ORF are identical, save for three indels in the upstream region. This close but not perfect match appears to have caused a split in the assembly graph that lead to the spurious contig. Additional information is in Supplemental Table S3. Again, there was no evidence for *pifA* in either the draft or ultimate assembly.

### FIGURES

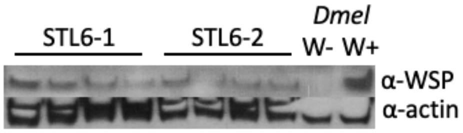


**Figure S1.** Western blot against *Wolbachia* Surface Protein (WSP). Controls are *Wolbachia*-infected and *Wolbachia*-uninfected *Drosophila melanogaster* ("*Dmel*").

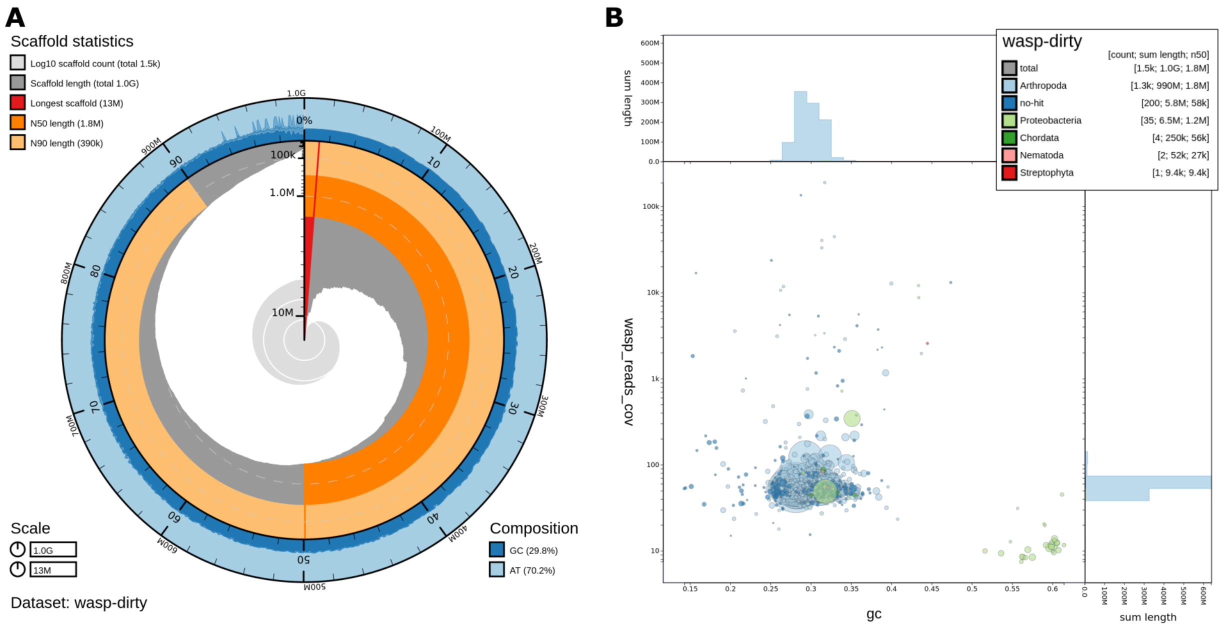


**Figure S2.** Draft assembly for *Leptopilina n. sp.* **(A)** snail plot with scaffold statistics **(B)** blobplot showing scaffold coverage, GC%, and taxonomy.
